# UV irradiation remodels the specificity landscape of transcription factors

**DOI:** 10.1101/2022.10.07.511343

**Authors:** Zachery Mielko, Yuning Zhang, Yiling Liu, Matthew A Schaich, Brittani Schnable, Debbie Burdinski, Sheera Adar, Miles Pufall, Bennett Van Houten, Raluca Gordan, Ariel Afek

## Abstract

Somatic mutations are highly enriched at transcription factor (TF) binding sites, with the strongest trend being observed for ultraviolet light (UV)-induced mutations in melanomas. One of the main mechanisms proposed for this hyper-mutation pattern is the inefficient repair of UV lesions within TF-binding sites, caused by competition between TFs bound to these lesions and the DNA repair proteins that must recognize the lesions to initiate repair. However, TF binding to UV-irradiated DNA is poorly characterized, and it is unclear whether TFs maintain specificity for their DNA sites after UV exposure. We developed UV-Bind, a high-throughput approach to investigate the impact of UV irradiation on protein-DNA binding specificity. We applied UV-Bind to ten TFs from eight structural families, and found that UV lesions significantly altered the DNA-binding preferences of all TFs tested. The main effect was a decrease in binding specificity, but the precise effects and their magnitude differ across factors. Importantly, we found that despite the overall reduction in DNA-binding specificity in the presence of UV lesions, TFs can still compete with repair proteins for lesion recognition, in a manner consistent with their specificity for UV-irradiated DNA. In addition, for a subset of TFs we identified a surprising but reproducible effect at certain non-consensus DNA sequences, where UV irradiation leads to a high increase in the level of TF binding. These changes in DNA-binding specificity after UV irradiation, at both consensus and non-consensus sites, have important implications for the regulatory and mutagenic roles of TFs in the cell.

## INTRODUCTION

Transcription factor (TF) proteins bind to specific loci in the genome to modulate gene expression. The recognition of specific, short DNA sites among the vast genomic background is highly affected by the chemical identities of the DNA bases (base recognition) as well as the local DNA structure (shape recognition) (1, 2). Any chemical reaction that modifies DNA bases or locally distorts the DNA can alter a TF’s binding affinity and specificity. Exposure to the sun’s ultraviolet (UV) rays promotes several types of DNA-distorting chemical reactions (**Fig. 1A**), with the most common being the formation of cyclobutane pyrimidine dimers (CPD) and 6-4 photoproducts (6-4PP) at pyrimidine dinucleotides (3). CPDs and 6-4PPs disrupt hydrogen bonding and distort the structure of the DNA by bending the phosphodiester backbone (4–6), which can affect both shape recognition and direct base readout by TFs. However, the impact of these UV-induced lesions on TF binding is poorly understood and has only been tested for a handful of DNA sites (7–11).

**Figure 1:**
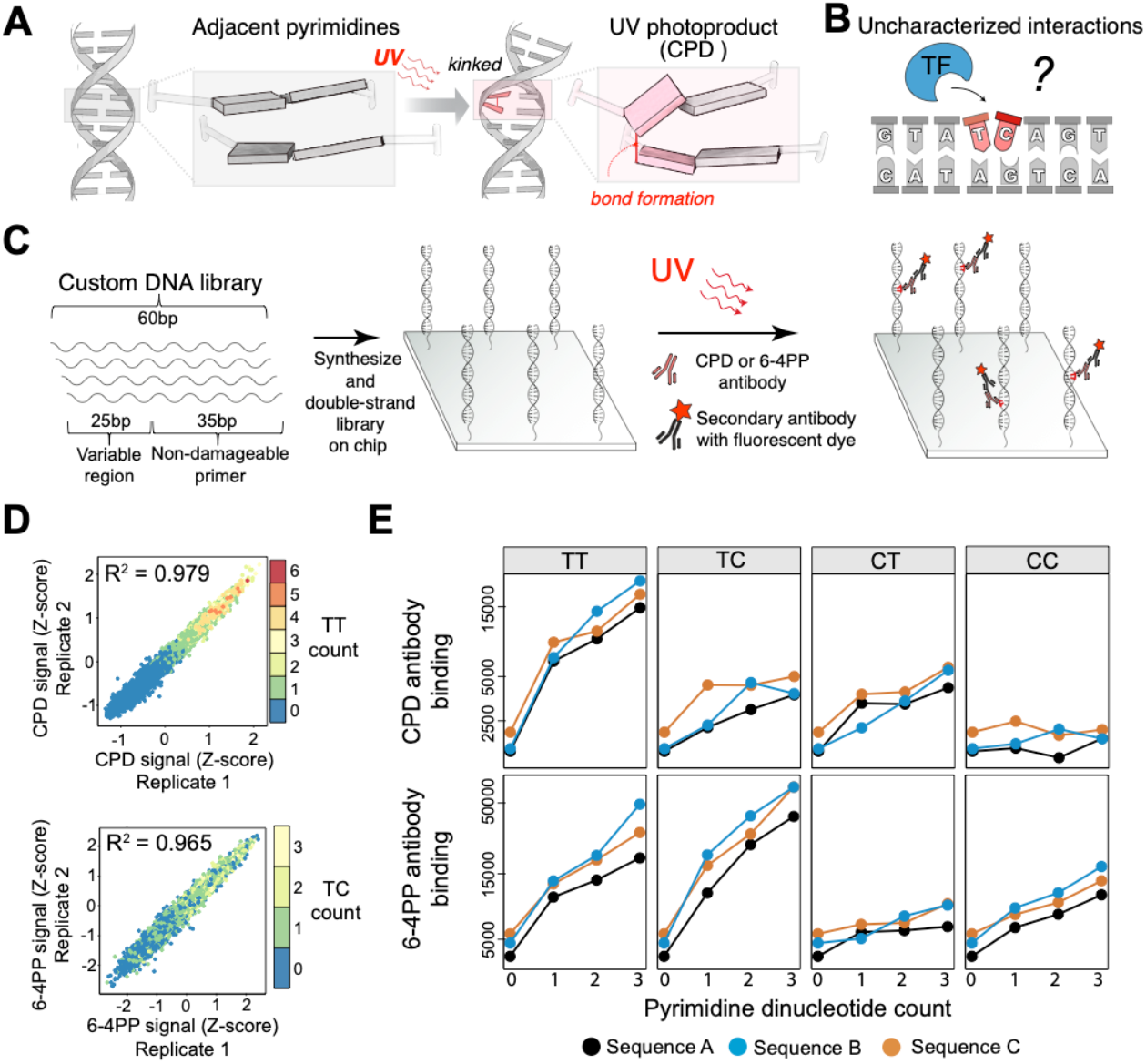
UV-Bind measures protein binding to UV-irradiated DNA. **(A)** UV light creates dipyrimidine photoproducts (CPD shown here), which distort the DNA. **(B)** These distortions are likely to affect TF-DNA recognition, which depends on both the sequence and the structure of the DNA. Binding of TFs to DNA sites containing UV-induced lesions is not well understood. **(C)** The UV-Bind methodology uses custom DNA libraries of up to 180,000 sequences printed on high-density DNA chips. The DNA chips are exposed to UV light, and photoproduct formation is measured using antibodies specific to the lesions of interest (here, CPD and 6-4PP). **(D)** CPD and 6-4PP antibody binding signals for all possible 7-mers, computed using a universal DNA library (Methods; **Table S1**). The R^2^ values represent the squared Pearson correlation coefficients between replicate experiments. Each data point is colored based on the number of TT or TC dinucleotides in each 7-mer, for CPD and 6-4PP, respectively. **(E)** CPD and 6-4PP antibody binding signal (median over 10-30 measurements; Methods) for three custom DNA sequences with an increasing number of pyrimidine dinucleotides (**Table S2**). The three sequences were selected so that the dipyrimidines were inserted in different sequence contexts. All plots show a significant increasing trend (Jonckheere’s trend test p-values between 0.01 and 10^−5^; **Table S2**) except for CC CPDs, which are expected to occur with very low frequency (19).

Understanding TF binding to UV lesions is critical for deciphering their hypothesized role in UV-induced mutagenesis, as well as the potential effects of UV irradiation on transcriptional regulation of gene expression. Specifically, recent genomic studies showed that nucleotide excision repair (NER) is less efficient at active TF binding sites, while mutation rates at these sites are significantly elevated, consistent with the hypothesis that TFs impair repair and increase mutagenesis at their binding sites (12, 13). Although the exact mechanisms by which specific TFs influence mutagenesis are not fully understood, there is growing evidence that TFs might compete with DNA repair proteins for the damaged DNA substrate, and thus effectively act as a barrier for the repair machinery (13–15). This competition, though, depends strongly on whether TFs can still bind efficiently to their target genomic DNA sites in the presence of UV lesions. The few, small-scale studies performed to date indicate that UV lesions typically reduce TF binding affinity, with decreases of up to 60-fold (7–11). However, the effects of UV irradiation on the binding preferences of any given TF are not predictable, and no high-throughput methods exist to characterize these preferences.

We present UV-Bind, an *in vitro* high-throughput technique for measuring protein binding to tens of thousands of DNA sequences containing UV-induced lesions. Our technique can be used with a variety of DNA library designs and DNA-binding proteins, including transcription factors, DNA repair proteins, and photoproduct-specific antibodies. When applied to ten TF proteins, UV-Bind revealed that UV lesions change the DNA recognition landscape of TFs, in a way that cannot be predicted from existing models of specificity for undamaged DNA. We also found that the TF binding preferences for UV-irradiated DNA, as characterized using UV-Bind, dictate the ability of TFs to compete with repair proteins (here, UV-DDB) that must recognize UV lesions to initiate repair. Thus, we expect our new technique and the findings in this study to lay the foundation for a comprehensive, mechanistic understanding of the role of TFs in DNA repair and mutagenesis in response to UV exposure.

## RESULTS

### UV-Bind provides quantitative measurements of protein binding to UV-irradiated DNA

UV-Bind is a new technique that uses high-density DNA arrays (chips) comprising of tens of thousands of DNA spots, with each spot containing millions of copies of a custom sequence of up to 60 base pairs (bp). We designed the DNA sequences to contain a 25bp variable region, followed by an undamageable 35bp “primer region” used to double-strand the DNA on the array (**Fig. 1C**). To ensure that the primer region is undamageable, we designed it to not contain any pyrimidine dinucleotides, nor purine dinucleotides (which would result in pyrimidine dinucleotides on the opposite strand). After double-stranding the DNA, we exposed the chip to a relatively large dose of UVC radiation (1.5 kJ/m^2^), as in previous studies of protein binding to UV-irradiated DNA (9, 11, 16). At the applied dose, UVC is known to create mostly CPD and 6-4PP lesions at pyrimidine dinucleotides (17, 18). The absolute frequencies with which each lesion forms at any particular pyrimidine dinucleotide in a particular sequence context are not known; however, at a broad level, the relative prevalence of CPDs is expected to be TT>TC>CT>>CC, while for 6-4PPs is TC>TT>CC>CT (19, 20).

To validate the formation of UV lesions in our assay, we incubated the irradiated chip with anti-CPD and anti-6-4PP antibodies, followed by secondary antibodies with Alexa dyes (**Fig. 1C**; Methods). Next, we scanned the DNA chip to measure the fluorescence intensity, which is directly proportional to the amount of antibody bound to the DNA at each spot. The antibody binding data was highly reproducible (R^2^ = 0.979 for CPD, 0.965 for 6-4PP; **Fig. 1D**; **Table S1**) and consistent with the expected prevalence of each dipyrimidine lesion type (19) (**Fig. 1D,E**). In addition, we found that the binding level for both antibodies increased with the addition of more dipyrimidines, as expected (**Fig. 1E, Table S2**).

### UV irradiation causes widespread changes in protein-DNA binding

To measure changes in protein-DNA binding specificity due to UV irradiation, we designed a custom DNA library covering all possible 7-mers. The library design is based on de Bruijn sequences (Supplementary Methods), similar to the well-established universal protein-binding microarray (uPBM) design (21). Standard 7-mer uPBM libraries contain 10,293 distinct 36bp sequences that together cover all possible 7-mers, with each 7-mer occurring in at least 16 distinct probes in the DNA library (21). We modified this design in order to reduce the average number of damageable dinucleotides per sequence (from 27 to six; **Fig. S1**). Our universal UV-Bind design uses shorter custom DNA probes (25bp), with each probe containing a variable 14bp “de Bruijn region” (named for the compact patterning used to cover the sequence space), flanked by 5 or 6 base pairs of undamageable sequence (Methods, **Fig. 2A, Fig. S1**). A total of 43,691 distinct 14-mers are sufficient to cover all possible 7-mers with at least 16 occurrences per 7-mer, thus providing comprehensive data on DNA-binding specificity.

**Figure 2:**
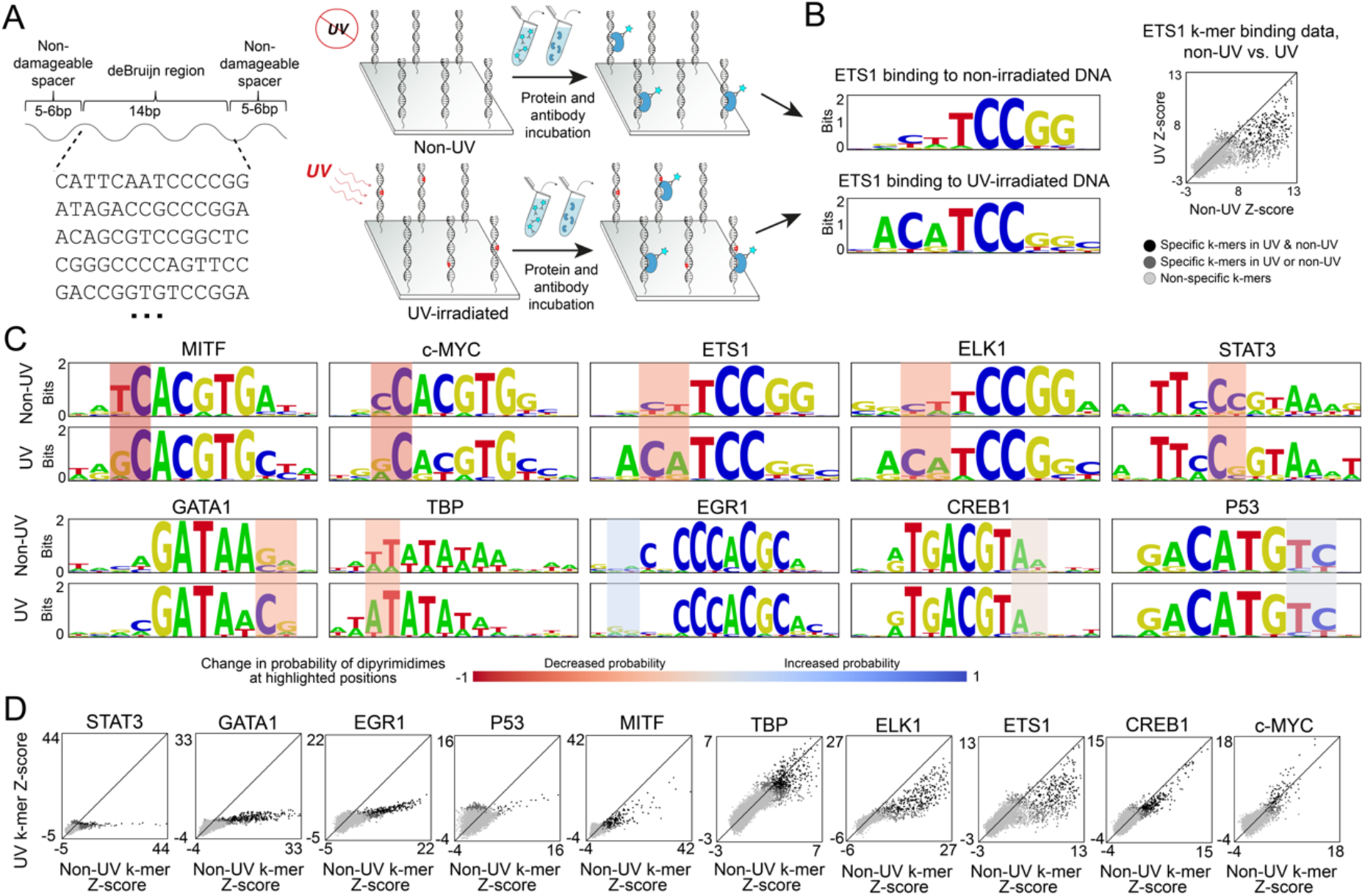
Comparison of TF binding specificities for UV-irradiated vs. non-irradiated DNA. **(A)** UV-Bind experiments using non-irradiated vs. UV-irradiated universal DNA libraries were used to comprehensively measure TF binding to all possible 7-mers. **(B)** TF binding specificities under the two conditions were compared using PWM binding models derived from the universal UV-Bind data (left), as well as the 7-mer binding intensity Z-scores (right). Data points are colored based on a binding enrichment score (E-score) cutoff of 0.35, which roughly corresponds to specific TF binding (22). **(C)** PWM comparisons in non-UV and UV conditions, across 10 TF proteins. For each TF, the dinucleotide position with the largest change in the probability of pyrimidine-pyrimidine (on either strand) is highlighted. **(D)** Binding Z-score comparisons between non-UV vs. UV conditions, for all 10 TF tested. Data points are colored as in panel B. TFs are ordered by the slope of the best fit line for the specific k-mers (black data points; **Table S3**).

We used UV-Bind with the modified universal DNA library to measure the effects of UV irradiation on the binding specificity of 10 TFs, selected to cover a variety of structural families. For each protein, we measured DNA binding to the UV-irradiated as well as the non-irradiated universal library (**Fig. 2A**), with high reproducibility (**Fig. S2**). From the fluorescence intensity data, we computed median intensities for all possible 7-mers (over the 16 or more probes containing each 7-mer), normalized binding scores for all 7-mers (Z-scores), and position weight matrices (PWMs), which provide an overview of the protein’s binding specificity for UV-irradiated vs. non-irradiated DNA (**Fig. 2B**).

Our data revealed that UV irradiation changes the DNA-binding specificity of all TFs tested, with the dominant effect being a decrease in binding to damageable sequences, i.e. to sequences that contain pyrimidine dinucleotides. This trend is illustrated in **Fig. 2C** through direct comparisons between PWM motif models generated from the non-irradiated vs. the UV-irradiated DNA libraries. For each dinucleotide position in the binding motifs, we calculated the change in the probability of having a pyrimidine-pyrimidine at that position, on either strand (**Fig. S3**). The positions with the largest change for each TF, which varies between −81% (for MITF) and +4% (for P53), are highlighted in **Fig. 2C**. For eight of ten TFs tested, the largest change corresponds to a decrease in the probability of pyrimidine dinucleotides, indicating that UV-induced damage at these positions is detrimental for TF binding.

For TFs that belong to the same structural family, we found that the changes in DNA-binding specificity after UV irradiation are similar. Basic helix-loop-helix proteins MITF and c-MYC have a reduced preference for pyrimidine dinucleotides in the immediate flank of the CACGTG core site, while ETS family members ETS1 and ELK1 both exhibit changes in specificity from **T**TCCGG to **A**TCCGG. We note that not all positions with high probabilities for pyrimidine dinucleotides showed a decrease in specificity. For example, the GA in GATA1’s **GA**TA core motif remains unchanged in the UV compared to the non-UV motifs, even though its reverse complement, TC, is damageable. However, UV irradiation reduced GATA1’s preference for pyrimidine dinucleotides at most positions around the GATA core, as shown in **Fig. S3A**. Over all 10 TFs tested, about two thirds of positions showed a decrease rather than an increase in the probability of pyrimidine dinucleotides, and the magnitude of the observed decreases was significantly larger than the magnitude of the observed increases (Mann-Whitney U test p-value = 0.002; **Fig. S3B**). This overall reduction in TF binding specificity after UV irradiation is also observed when comparing the k-mer binding data between the non-UV and UV conditions (**Fig 2D**; **Table S3**).

### The magnitude of TF binding changes due to UV irradiation varies widely depending on the precise binding site sequences and the positions of UV lesions

To further investigate the effects of UV irradiation on TF-DNA binding specificity, we performed UV-Bind experiments with DNA libraries focused on specific binding sites and single or double nucleotide variations in these sites. This library design allows us to investigate effects of UV lesions in binding sites that do not necessarily match the consensus represented by the PWMs. Furthermore, by starting with non-damageable DNA binding sites, we can more easily interpret the UV-Bind data and pinpoint the effects of UV lesions at specific positions. For these analyses, we first focused on the TF MITF and its canonical E-box site, CACGTG (see **Fig. S4A** for analyses of an alternative MITF core binding site, the M-box CACATG). MITF is a critical regulatory factor in melanocytes, where it is involved in cell proliferation, response to UV irradiation, and DNA damage repair (23). According to our universal UV-Bind data, MITF showed the largest decrease in the probability of pyrimidine-pyrimidine across all dinucleotide positions and all TFs tested (**Fig. 2C**). Focusing on this dinucleotide position at the flank of the E-box site, we measured the change in MITF-DNA binding due to UV irradiation for all N<u>TC</u>ACGT<u>GA</u>N and N<u>CC</u>ACGT<u>GG</u>N sites (underlining marks the dinucleotide position of interest, on both sides of the palindromic E-box). To assess the significance of the observed changes, we included in the DNA library a set of 447 non-damageable DNA sequences with a wide range of binding affinities for MITF (Supplementary Methods). We used these control sequences to fit an ordinary least squares (OLS) model and we computed the 99% prediction interval that reflects the experimental variation in MITF binding intensity expected at DNA sites not affected by UV irradiation (**Fig. 3A**, dotted lines). We then used this interval to assess whether MITF binding to damageable N<u>TC</u>ACGT<u>GA</u>N and N<u>CC</u>ACGT<u>GG</u>N sites is significantly affected by UV-induced lesions.

**Figure 3.**
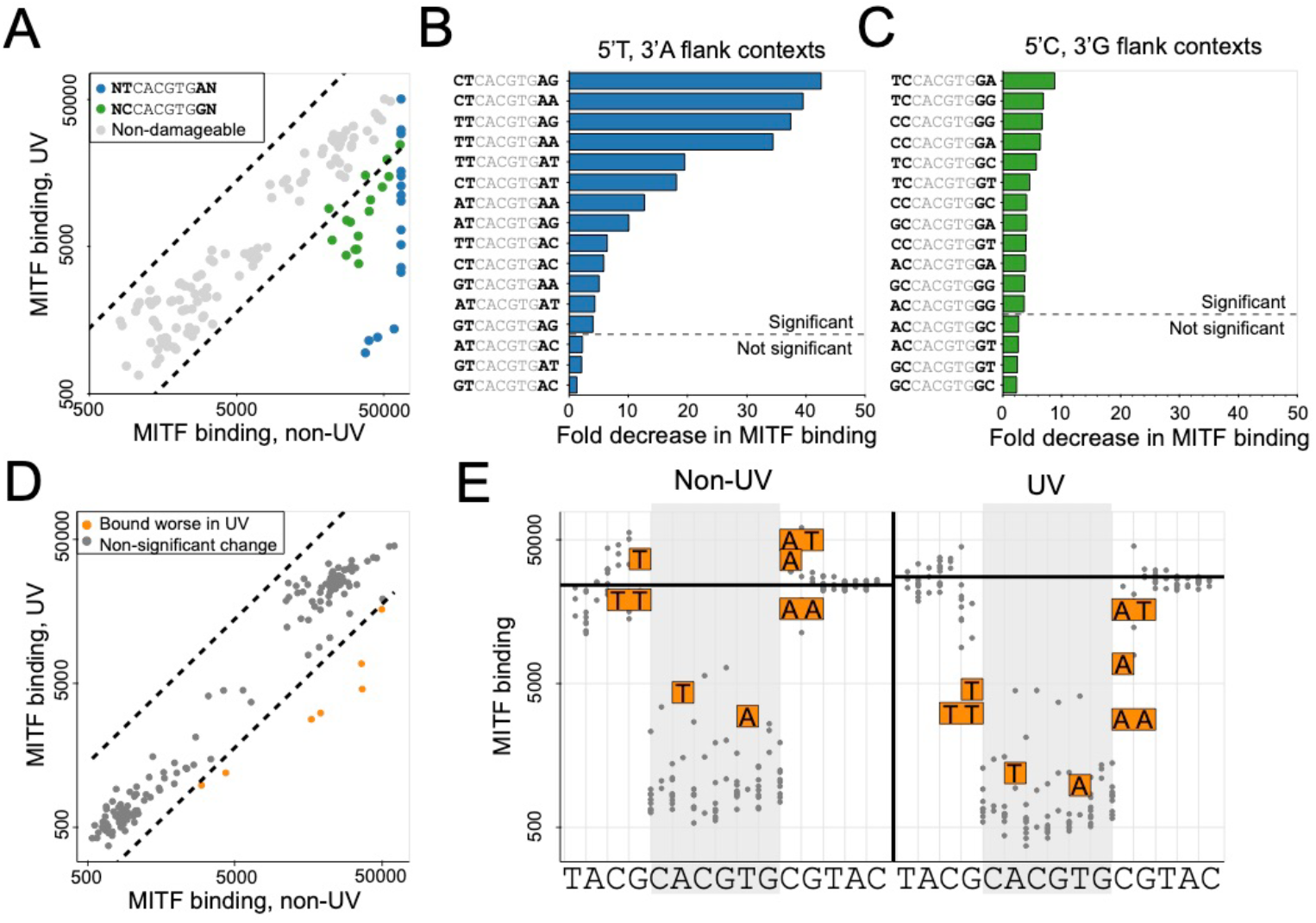
UV-Bind reveals a wide range of TF binding changes due to UV irradiation, beyond what is captured by PWM models. **(A)** MITF binding levels in UV vs. non-UV conditions (i.e. to UV-irradiated vs. non-irradiated DNA), for binding sites that vary in their immediate flanking sequence. Blue: <u>NT</u>CACGTG<u>AN</u> sites. Green: <u>NC</u>CACGTG<u>GN</u> sites. Grey: non-damageable sites. Dotted lines mark the 99% prediction interval (Methods) computed from non-damageable sites. **(B)** Fold decrease in MITF binding, computed between UV and non-UV conditions, for all <u>NT</u>CACGTG<u>AN</u> sites. Sites are ranked by the fold decrease from non-UV. Non-significant changes (i.e. changes that fall within the 99% prediction interval computed from non-damageable sequences) are shown below the dashed line. **(C)** Similar to panel B, but for all <u>NC</u>CACGTG<u>GN</u> sites. **(D)** MITF binding levels in UV vs. non-UV conditions, for a DNA library that contains all mononucleotide and dinucleotide variants of DNA sequence TACG<u>CACGTG</u>CGTAC (E-box site is underlined). Dotted lines mark the 99% prediction interval (Methods) computed from the non-damageable sites. All variants that fall within this interval are shown in grey. All variants that fall outside the interval are shown in orange. **(E)** Same data as in panel D, but shown as saturation variation plots relative to the wild-type reference sequence. Each grey point represents one of the grey variants from panel D, which did not show a significant difference in MITF binding between UV and non-UV. Letters with orange boxes around them show the variations that led to significant differences between non-UV and UV conditions, and correspond to the orange data points in panel D. The black horizontal line indicates the binding signal of the reference sequence. Grey shading highlights the core CACGTG region. All measurements in the figure are median values over 20 replicates (**Table S4**).

We found that UV irradiation can cause a wide range of decreases in MITF binding to sites containing the preferred 5’ T and 3’ A flanks (**Fig. 3B**), from a 42-fold decrease at <u>CT</u>CACGTG<u>AG</u> to a 1.3-fold decrease at <u>GT</u>CACGTG<u>AC</u>. Our data also revealed a trend not visible in the MITF PWMs: having CT or TT pyrimidine dinucleotides immediately flanking CACGTG has the largest effects on MITF binding following UV irradiation, indicating that UV lesions at these positions are highly detrimental to binding. Interestingly, for the less preferred MITF sites NCCACGTGGN (**Fig. 3C**), the decrease in binding after UV irradiation was less pronounced, between 8.8-fold and 2.3-fold, possibly due to the lower propensity of UV damage at CC dinucleotides. Similar to the trend observed for NTCACGTGAN sites, the presence of pyrimidine dinucleotides immediately flanking the CACGTG core had the largest effects on MITF binding. Overall, our data showed that UV lesions in the immediate flanks of the CACGTG sites, as well as lesions that form between flanks and the core are detrimental to MITF binding, with the magnitude of the effects depending on the precise binding site sequence.

We also used UV-Bind to measure the effects of UV irradiation on MITF binding to all mononucleotides and dinucleotide variants in its binding site and flanking regions (**Fig. 3D,E**). For this DNA library design, we started with a site that was completely non-damageable, and thus the immediate flanks are different from the ones analyzed above. For this particular MITF site and its variants, we found that UV irradiation resulted in small, mostly non-significant binding changes, as most sequences tested fell within the 99% prediction interval computed from non-damageable sequences (**Fig. 3D**, grey data points). Of the 172 sequences tested, only 7 (**Fig. 3D**, orange data points) showed a significant binding difference between UV and non-UV. Five of these sequences were variants that created a CT or TT dinucleotide immediately upstream or downstream of the E-box CACGTG; these variants were bound similarly to the wild-type site in the absence of UV irradiation (**Fig. 3E**, left), but they were bound poorly compared to the non-damageable wild-type site when tested after UV irradiation (**Fig. 3E**, right). These results are fully consistent with those in **Fig. 3A-C**, showing that CT or TT dinucleotides immediately upstream or downstream of the E-box lead to significant decreases in MITF binding after UV exposure. In addition to confirming this trend, our data also revealed two E-box variants that are significantly affected by UV: C<u>T</u>CGTG and its reverse complement site CACG<u>A</u>G (**Fig. 3E**; variants highlighted in orange within the core CACGTG region). While all other nucleotide variations in the E-box region led to similar decreases in MITF binding to non-irradiated vs. irradiated DNA, the two variants mentioned above, which result in the formation of damageable dipyrimidines CT and TC, show a more significant decrease for the UV-irradiated DNA (**Fig. 3E**, right) compared to the non-irradiated DNA (**Fig. 3E**, left), indicating that UV lesions in the core E-box site are detrimental to MITF binding. This trend was not observed in the PWM-based analysis, which is not surprising given the MITF sites with damageable dinucleotides in the core are weakly bound and thus not well captured by the PWMs. Additional analyses illustrating the complexity of the effects of UV irradiation on TF-DNA binding specificity are presented in **Fig. S4**, based on UV-Bind data for the MITF M-box site and for DNA binding sites of CREB1 and EGR1 (**Table S4**).

### UV irradiation leads to high levels of TF binding at sequences lacking consensus binding sites

In addition to the DNA sequences described above, our UV-Bind libraries included control sequences containing sites from protein-DNA structures available in the Protein Data Bank (PDB) (24) for a wide range of TFs, as well as single nucleotide variants of these PDB sites (Supplemental Methods). Surprisingly, when analyzing our UV-Bind data we found that control sequences containing a binding site for the TF RELA (a protein not tested in our study) and its variants showed highly increased binding by TFs CREB1 and EGR1 after UV irradiation (**Fig. S5A,B**). The RELA consensus binding site is very different from the consensus sites of CREB1 and EGR1. In addition, an in-depth analysis of these control sequences using non-UV k-mer binding data for CREB1 and EGR1 confirmed that they did not contain k-mers with high binding enrichment scores, i.e. E-scores, which is typical of consensus sites (**Fig. S5C-F**). A closer investigation, though, revealed that the control sequences of interest were composed of overlapping k-mers for which the binding E-scores were significantly higher in the UV versus the non-UV conditions (**Fig. S5D,F**). We note that these individual E-scores were still generally lower than what we expect for consensus binding sites, which oftentimes show multiple consecutive k-mers with E-score > 0.35 (**Fig. S5C,E**). Nevertheless, the k-mers showed a large increase in their E-scores in UV compared to non-UV conditions, leading us to hypothesize that by concatenating such ‘UV-preferred’ k-mers we can generate DNA sequences that are strong binders *after* UV-irradiation, even though they lack consensus binding sites for the tested TFs.

We tested this hypothesis by creating new DNA libraries for CREB1 and EGR1 containing sequences generated by concatenating ‘UV-preferred’ k-mers. To identify such k-mers we first trained an OLS model on replicate non-UV measurements of k-mer E-scores, in order to capture the experimental variability in these binding scores (Supplementary Methods, **Fig. S6B**). For this analysis we focused on k-mer E-scores, rather than Z-scores or median binding intensities. E-scores, which are widely used in analyses of universal PBM data (25–28), are computed using a rank-based, nonparametric statistic based on a modified form of the Wilcoxon-Mann-Whitney test (21). Consequently, they are robust to outliers and are comparable between experiments for different proteins or different experimental conditions (29). Here, we compared E-scores for TF binding to UV-irradiated vs. non-irradiated DNA. We used an OLS model derived from non-UV E-score data to identify k-mers that significantly change due to UV irradiation, i.e. they fall outside the 99% prediction interval derived from replicate non-UV measurements (Supplementary Methods; **Fig. S6A,B**). Next, we computationally generated new DNA sequences by concatenating overlapping ‘UV-preferred’ k-mers (**Fig. S6C**). To avoid testing highly similar sites, we clustered the generated sequences using affinity propagation (30, 31) and chose cluster representatives based on the maximum sum of preference scores (Methods), for a total of 3,200 unique sequences for CREB1 and 1,884 unique sequences for EGR1 (**Fig. S6D,E**). Indeed, our UV-Bind data for CREB1 and EGR1 using these newly generated DNA libraries confirmed that by concatenating ‘UV-preferred’ k-mers we obtain sequences that have a much higher level of TF binding to UV-irradiated compared to non-irradiated DNA (**Fig. 4**, black data points and distributions; **Table S5**).

**Figure 4:**
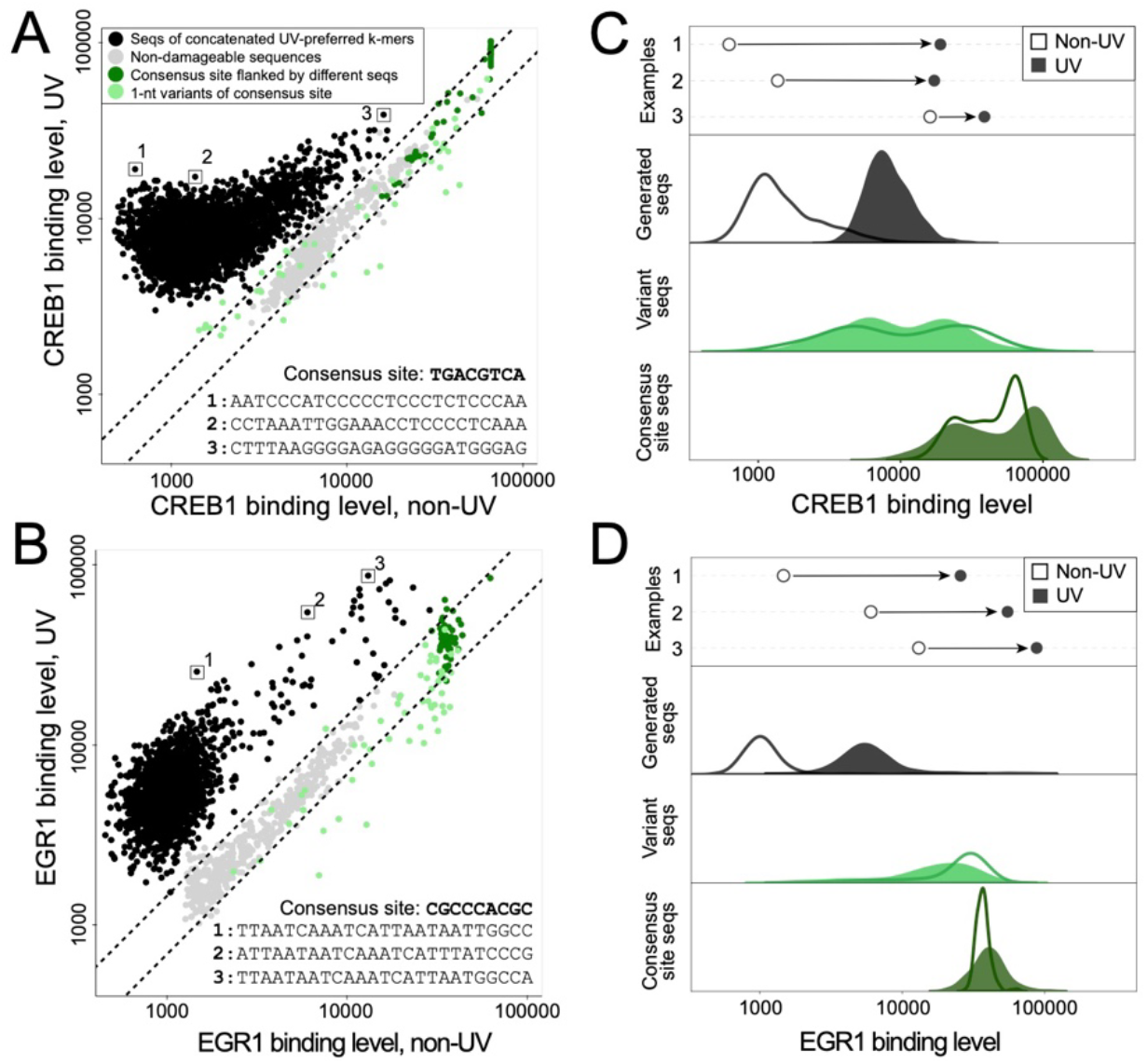
UV irradiation leads to high levels of TF binding at certain DNA sequences that lack consensus binding sites. **(A,B)** Binding of CREB1 and EGR1 to UV-irradiated versus non-irradiated DNA (Y-axes and X-axes, respectively). Grey points: sequences without any pyrimidine dinucleotides, which were used to train an ordinary least squares regression model (Methods). The model was then used to generate a 99% prediction interval of these sequences, marked by the dashed lines, which can be interpreted as follows: if UV irradiation does not affect TF binding specificity, then we would expect 99% of the tested sequences to fall within this interval. Dark green points: sequences containing high-affinity consensus sites for the tested TFs, embedded within different flanking sequences. The precise consensus site sequences are shown. Light green points: single-nucleotide variants of the consensus binding sites for CREB1 and EGR1. As expected, these variant sites show a wide range of TF binding levels. Black points: sequences generated by concatenating ‘UV-preferred’ k-mers. Data points represent medians over 20 replicate binding measurements for grey, green, and light green groups, and 3 replicates for the black group. (**C,D**) Comparisons between the distributions of TF binding levels for UV-irradiated and non-irradiated DNA. The groups of sequences are the same as in panels A and B. Distributions shown are gaussian kernel density estimates of the scatterplot data in panels A and B. Arrows indicate the change from non-UV to UV for three representative sequences.

### ETS1 competes with UV-DDB in a manner consistent with its binding specificity for UV-irradiated DNA

Next, we asked whether TF binding to UV-irradiated DNA can interfere with the recognition of UV-induced lesions by DNA repair proteins. To answer this question, we used UV-Bind to measure the DNA-binding activity of the UV-DDB (UV-damaged DNA-binding protein) complex, which plays a major role in global genomic nucleotide excision repair as the initial sensor for UV-induced photoproducts (32). First, we verified the binding activity of recombinant UV-DDB (here, a complex of His-tagged DDB2 and Flag-tagged DDB1 (33); Methods) in our chip-based assay by leveraging the DNA library used to measure damage formation using the anti-CPD and anti-6-4PP antibodies (**Fig. 1E**). As expected, we found that UV-DDB binding measurements to UV-irradiated DNA are highly reproducible, and that UV-DDB binding signal increases with an increasing number of pyrimidine dinucleotides (**Fig 5A-C**; **Table S2**). Our high-throughput data also shows trends in the binding specificity of UV-DDB: TC photoproducts show the highest level of binding, followed by TT and CC (whose ranking depends on the sequence context) and then by CT photoproducts, which show the lowest level of binding (**Fig. 5C**). K-mer binding data, which shows similar trends, and the UV-DDB DNA-binding motif derived from the k-mer data are available in **Table S6**. When tested against non-irradiated DNA, UV-DDB showed very low signal and no significant trend for sequences with an increasing number of dipyrimidines (**Table S2**), confirming its expected specificity for UV-induced photoproducts.

**Figure 5:**
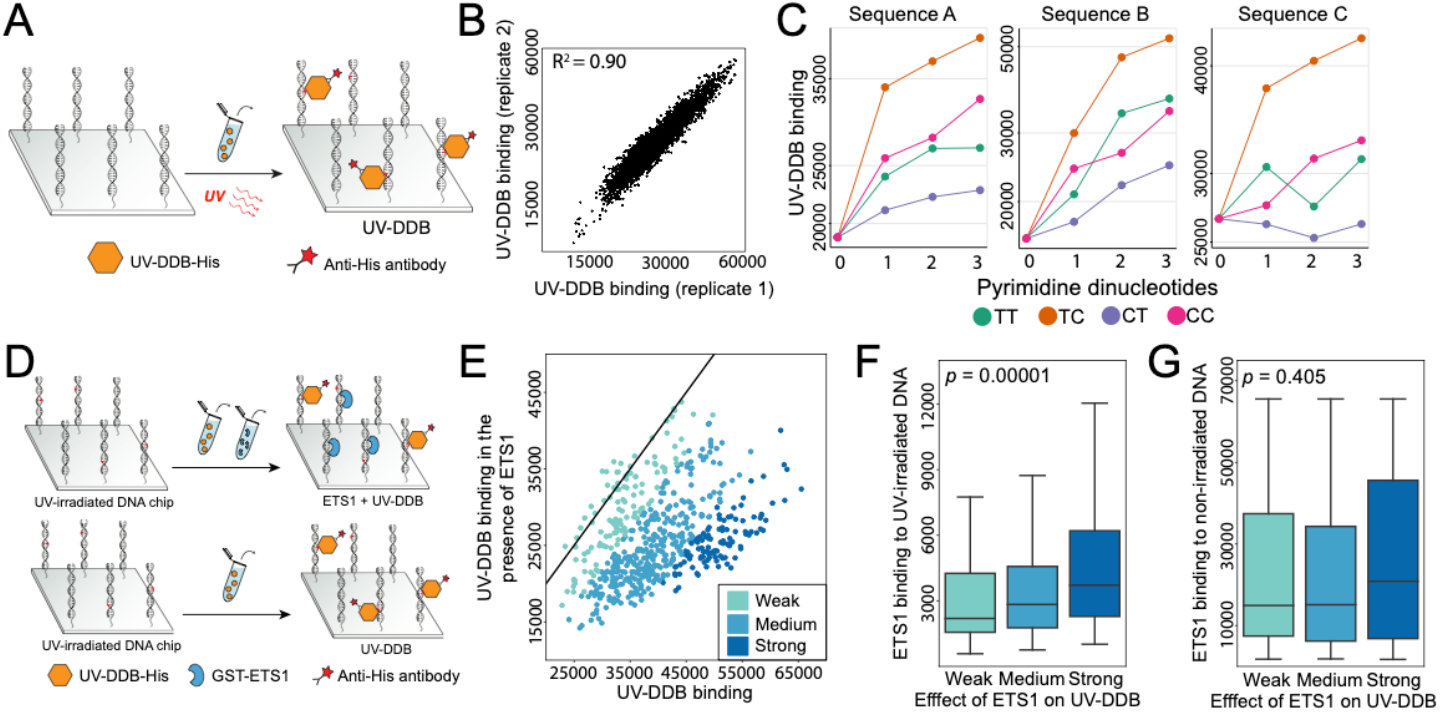
Transcription factor EST1 competes with repair protein UV-DDB for binding to UV-irradiated DNA. **(A)** UV-Bind was used the measure UV-DDB binding to UV-irradiated DNA in a high-throughput manner. **(B)** UV-DDB binding data is highly reproducible (R^2^ = 0.90). Measurements shown are median values among 10-30 replicate spots. **(C)** UV-DDB binding levels at UV-irradiated sequences containing an increasing number of dipyrimidines. One, two, or three dipyrimidines were inserted in three different sequence contexts (**Table S2**). For most dipyrimidines and sequence contexts, we found a significant increasing trend in UV-DDB binding (Jonckheere’s trend test p-value < 0.05); the only exceptions were TT and CT in the third sequence context tested, which was the most AT-rich (**Table S2**). Measurements shown are median values among 10-30 replicate spots. **(D)** Overview of the UV-DDB-ETS1 competition experiment. UV-irradiated chambers with the same library design were compared, one chamber with UV-DDB alone and another with both ETS1 and UV-DDB added at the same time. The concentration of UV-DDB was kept constant (Supplementary Methods). **(E)** UV-DDB-ETS1 competition data for the ETS1 binding site in crystal structure PDB ID: 2NNY, and all its single nucleotide variants. Sequences are grouped into three bins colored light to dark based on the distance from the diagonal (black line), indicating the strength of the ETS1 effect on UV-DDB. **(F)** The ETS1 binding level for UV-irradiated DNA correlates with the strength of the ETS1 effect on UV-DDB (Jonckheere’s trend test p-value = 10^−5^). **(G)** The ETS1 binding level for non-irradiated DNA does not show a significant correlation with the strength of the ETS1 effect on UV-DDB (Jonckheere’s trend test p-value = 0.405). Similar trends were seen across different sequences, binning methods and statistical tests (**Fig. S7C,D Table S7**, Supplementary Methods).

Having verified the DNA-binding activity of UV-DDB in UV-Bind assays, we next measured the effect of transcription factor ETS1 on the binding level of UV-DDB for UV-irradiated DNA, focusing on two subgroups of DNA sequences derived from high-affinity consensus binding sites for the ETS1 protein. As mentioned above, our UV-Bind libraries included control sequences containing TF-binding sites from co-crystal structures available in PDB (24). Two of these sites (PDB IDs 2NNY and 1DUX; Supplementary Methods) have high binding affinity for ETS1 (**Fig. S7A**), and they were included together with all their single nucleotide variants in our UV-Bind library. To test the competitive effect of ETS1 on UV-DDB, we incubated the UV-irradiated DNA library with both proteins simultaneously, we measured the level of UV-DDB binding, and compared it to the binding level of UV-DDB in the absence of ETS1, keeping the concentration of the repair protein constant (**Fig. 5D**; Supplementary Methods). We found that the UV-DDB binding level decreases in the presence of ETS1, consistent with the TF competing with the repair protein for DNA binding (**Fig. 5E**, **Fig. S7B, Table S7**). Next, we grouped the tested sequences into three bins based on whether ETS1 had a strong, medium, or weak effect on UV-DDB binding, reflected by the distance from the diagonal. We found that the magnitude of the competitive effect of ETS1 on UV-DDB correlated with the specificity of the TF for UV-irradiated DNA (**Fig. 5F**; Jonckheere test p-value = 10^−5^), indicating that the reduction in UV-DDB binding level is due to the TF’s specificity. When using the binding specificity of ETS1 for non-irradiated DNA, we did not find a significant correlation with the effect of ETS1 on UV-DDB (**Fig 5G**; Jonckheere test p-value = 0.405). Similar results were obtained at the second ETS1 site tested (**Fig. S7B-D**), and these results were robust with respect to the binning method and the statistical test used to determine if there is a significant increasing trend correlating ETS1 binding specificity with the effect of ETS1 on UV-DDB (**Table S7**). Overall, these results show that the ETS1 binding specificity for UV-irradiated DNA is needed to explain the TF’s competition with UV-DDB. More broadly, our analyses point to the importance of characterizing the specificity of TFs for UV-damaged DNA as a necessary step in our efforts to understand how TFs compete with repair proteins, reduce the efficiency of UV lesion repair, and potentially promote UV-induced mutagenesis at TF binding sites.

## DISCUSSION

UV irradiation causes changes in DNA-binding across multiple transcription factors and binding contexts. Using our new UV-Bind technique, we identified changes in binding specificity for all ten TFs tested, and we found that binding to canonical sites is mostly negatively impacted by UV irradiation (**Fig. 2**). These results greatly expand upon previous studies (7, 8, 10, 11), which only tested TF binding to a small number of sequences with synthesized TT CPD lesions. For example, Tommasi *et al*. (7) previously found that UV lesions can cause reductions in TF binding ranging between 11-fold and 60-fold. Our study confirms that effects of this magnitude do occur (**Fig. 3**) but, depending on the TF, binding site, and precise location of the lesion, the effects of UV-irradiation on TF-DNA binding can vary widely, with some lesions even having the potential to slightly increase binding (**Fig. S4**). Overall, our results demonstrate the need for high-throughput measurements of TF binding to UV-irradiated DNA in order to fully understand how UV lesions affect TF-DNA recognition.

Surprisingly, our UV-Bind data also showed increased levels of TFs bound to DNA sequences lacking any resemblance to the TFs’ canonical sites (**Fig. 4, Fig. S6**). Although additional studies are needed to understand this phenomenon and its potential effects in the cell, our finding is consistent with a previous report showing that UV-irradiated non-canonical DNA sites can act as decoys to titrate TFs away from their regulatory sites and thus affect gene expression (9). Specifically, Vichi *et al*. reported that increased UV doses led to increased TBP binding to a sequence without TATA elements, and hypothesized that the TF might be sequestered away from its canonical site, causing a reduction in gene transcription in fibroblasts (9). Additional TBP inserted into the cells prior to UV irradiation partially rescued the reduction in transcription, consistent with their hypothesis. Our results show that the effect of UV irradiation on TF binding to sequences without canonical sites extends to CREB1 and EGR1. Furthermore, UV-Bind data can be used to predict DNA sequences where this effect is likely to occur (**Fig. 4**). For TFs such as CREB1, which binds to approximately 17% of promoters in the human genome, decoy non-canonical sites caused by UV irradiation could have a widespread impact on gene expression (34).

Our universal UV-Bind data showed that the main effect of UV irradiation was a decrease in TF binding specificity. Given this finding, we asked whether TFs bound to UV-irradiated DNA would be able to compete with repair proteins that must recognize UV lesions and initiate their repair, as hypothesized in the literature (12, 13). This question is particularly relevant for transcription factors from the ETS family (including ETS1 and ELK1 tested here), for which there are conflicting reports regarding the mechanisms by which these TFs increase mutagenesis at their binding sites in melanoma. Several studies showed that ETS binding sites in the human genome are hotspots for CPD formation, with ETS binding favoring the formation of these lesions due to TF-induced structural changes in the binding sites (16, 35, 36); the effect of ETS binding on the formation of 6-4PP lesions is currently unknown. Other studies, though, reported a reduced rate of DNA repair at ETS binding sites, pointing to interference with repair as a contributing factor to the hyper-mutation pattern observed at these sites (12). Our UV-Bind data showed that UV irradiation causes a pronounced reduction in the DNA-binding specificity of both ETS factors tested, ETS1 and ELK1 (**Fig. 2**), which raises doubts about the potential of ETS factors to compete with DNA repair. UV-Bind allowed us to directly test this competition (**Fig. 5**). We found that despite the overall reduction in ETS1-DNA binding specificity in the presence of UV lesions, the TF can still compete with UV-DDB, a major repair protein responsible for the recognition of UV lesions and the initiation of repair through the global genomic NER pathway. Furthermore, we show that competition with UV-DDB is better explained by the ETS1 specificity for UV-irradiated DNA, compared to the canonical non-irradiated DNA specificity (**Fig. 5F,G**). Additional studies are needed to dissect, at high resolution, which mutations in ETS binding sites can be enhanced in living cells through each mechanism, i.e. DNA-bound ETS factors promoting the formation of UV lesions vs. ETS factors competing with repair proteins for binding at UV-damaged sites. Nevertheless, our study shows that competition between TFs and DNA repair proteins can occur even when UV irradiation reduces the TF’s DNA-binding specificity, supporting a role for TFs in UV-induced mutagenesis through interference with DNA repair.

## METHODS

### UV-Bind Assay

High-density single-stranded DNA arrays were purchased from Agilent, in either 8×60k or 4×180k formats (8 or 4 identical chambers, with approximately 60,000 or 180,000 DNA spots in each chamber, respectively). The DNA libraries synthesized on the arrays contained de Bruijn sequences covering all 9-mers, TF binding sites of interest and variants of these sites, and/or control sites, depending on the precise experiment (see Supplementary Methods for details). The DNA was double-stranded on the arrays by primer extension, as described in prior work (2). In order to irradiate only certain parts of the array, and protect the rest from UV irradiation, a cover/gasket slide was cut to the desired size are used to partially cover the DNA array. Next, the gasket slide was fully covered using Cryo-Babies® labels, and the array-gasket slide sandwich was placed in an open container with 1x PBS. To irradiate the DNA on the array, we used a Stratalinker® UV Crosslinker 1800 instrument and conducted a series of irradiations with UVC. Using the energy mode, the container was irradiated with 100 J/m^2^, randomly repositioned inside the crosslinker 15 times to achieve 1,500 J/m^2^. Midway through the series, the PBS was replaced in order to keep the solution at room temperature. This procedure resulted in a DNA array with some chambers of the array irradiated by UV and others not irradiated. Finally, the protein and antibody binding steps were performed as in standard PBM assays (29), and the fluorescent signal of bound protein for each DNA spot was determined using a GenePix® 4400A microarray scanner and the GenePix® Pro 7.3 software.

### Protein Expression and Purification

Full-length human ETS1, ELK1, GATA1, MYC (c-MYC), MITF and UV-DDB were expressed and purified as described in previous literature, with either a His_6_ or a GST tag (2, 33, 37). DNA-binding domains for human EGR1 (residues 335-423), phosphorylated STAT3 (residues 128-715), and *A.thaliana* TBP were purified and expressed as described previously (38–40). Full length human GST-tagged P53 protein was purchased from Abcam (Catalog number: ab43615), and His_6_-CREB1 was purchased from Origene (Catalog number: TP760318). Primary antibodies for CPD and 6-4PP photoproducts were purchased from Cosmo Bio USA (Catalog numbers: CAC-NM-DND-001 and CAC-NM-DND-002 respectively); these antibodies were first described by Mori et al. (41), who verified their specificity for CPD and 6-4PP photoproducts in both single-stranded and double-stranded DNA.

The following anti-His fluorophore-conjugated antibodies were purchased from Qiagen and used in UV-Bind assays for His-tagged proteins: Penta-His Alexa488-conjugated antibody and Penta-His Alexa647-conjugated antibody (Catalog numbers: 35310 and 35370 respectively). Anti-GST Alexa488-conjugated antibody was purchased from ThermoFisher Scientific (Catalog number: A11131) and Anti-GST Alexa647-conjugated antibody was purchased from Cell Signaling Technology (Catalog number 3445S). Anti-mouse Alexa488- and Alexa647-labelled antibodies (Cell signaling technology, Catalog numbers 2350S and 3445S) were used as secondary antibodies for anti-CPD and anti-6-4PP experiments.

### Data Processing and Analysis

The raw data collected using the GenePix® 4400A microarray scanner was processed using the Seed and Wobble suite (21), with modifications to adapt the k-mer data to our UV-Bind universal design, as described in Supplementary Methods. For each unique sequence tested, median values over the spots containing that sequence were used unless otherwise noted (i.e. for the universal design sequences and for the competition tests). The number of replicate spots for each sequence varied between 3 and 20, depending on the DNA library.

For comparisons between non-UV and UV conditions, a scaling transformation was applied based on sequences in the library without pyrimidine dinucleotides (Supplementary Materials) in order to account for effects in fluorescence intensity that were not related to binding of the tested protein to UV lesions. TF-specific OLS models were trained on all sequences without pyrimidine dinucleotides, to capture the observed variation in fluorescence at sequences that could not form pyrimidine photoproducts. Measurements were categorized as significantly or not significantly different than expected based on a 99% prediction interval of the model. One assumption of OLS models is that the data is homoskedastic (constant variation in the data), which is not an assumption that can be made for UV-Bind data. For this reason, our OLS models used heteroskedasticity-consistent (HC) standard errors, and specifically the HC3 standard errors (42, 43).

Comparisons for increasing trends observed for anti-CPD and anti-6-4PP antibodies, as well as for UV-DDB, at sequences with an increased number of dipyrimidine were performed using a Jonckheere trend test with an alternative hypothesis of an increasing trend and 10^5^ permutations. A similar approach was used to evaluate the significance on increasing trends observed in the competition data. Additional details can be found in the Supplementary Materials.

## ACKNOWLEDGEMENTS

The authors would like to thank Hana Wasserman, Harshit Sahay and Wei Zhu for thoughtful comments and discussion on this manuscript. Funding to support this work was provided by the US-Israel Binational Science Foundation grant BSF-2019272 (R.G. and S.A.), the Duke School of Medicine Precision Genomics Collaboratory-OBGE Graduate Student Pilot Research Grant (Z.M.) and NIH grants R01-GM135658 (R.G.) and R35ES031638 (B.V.H.).

## REFERENCES

1. M. Slattery et al., Absence of a simple code: how transcription factors read the genome. Trends Biochem Sci 39, 381–399 (2014).

2. A. Afek et al., DNA mismatches reveal conformational penalties in protein-DNA recognition. Nature 587, 291–296 (2020).

3. J. Cadet, T. Douki, Formation of UV-induced DNA damage contributing to skin cancer development. Photochem Photobiol Sci 17, 1816–1841 (2018).

4. J. K. Kim, D. Patel, B. S. Choi, Contrasting structural impacts induced by cis-syn cyclobutane dimer and (6-4) adduct in DNA duplex decamers: implication in mutagenesis and repair activity. Photochem Photobiol 62, 44–50 (1995).

5. H. Park et al., Crystal structure of a DNA decamer containing a cis-syn thymine dimer. Proc Natl Acad Sci U S A 99, 15965–15970 (2002).

6. R. P. Rastogi,Richa, A. Kumar, M. B. Tyagi, R. P. Sinha, Molecular mechanisms of ultraviolet radiation-induced DNA damage and repair. J Nucleic Acids 2010, 592980 (2010).

7. S. Tommasi, P. M. Swiderski, Y. Tu, B. E. Kaplan, G. P. Pfeifer, Inhibition of transcription factor binding by ultraviolet-induced pyrimidine dimers. Biochemistry 35, 15693–15703 (1996).

8. X. Liu, A. Conconi, M. J. Smerdon, Strand-specific modulation of UV photoproducts in 5S rDNA by TFIIIA binding and their effect on TFIIIA complex formation. Biochemistry 36, 13710–13717 (1997).

9. P. Vichi et al., Cisplatin- and UV-damaged DNA lure the basal transcription factor TFIID/TBP. EMBO J 16, 7444–7456 (1997).

10. Y. Kwon, M. J. Smerdon, Binding of zinc finger protein transcription factor IIIA to its cognate DNA sequence with single UV photoproducts at specific sites and its effect on DNA repair. J Biol Chem 278, 45451–45459 (2003).

11. S. Sivapragasam et al., CTCF binding modulates UV damage formation to promote mutation hot spots in melanoma. EMBO J 40, e107795 (2021).

12. R. Sabarinathan, L. Mularoni, J. Deu-Pons, A. Gonzalez-Perez, N. Lopez-Bigas, Nucleotide excision repair is impaired by binding of transcription factors to DNA. Nature 532, 264–267 (2016).

13. J. Frigola, R. Sabarinathan, A. Gonzalez-Perez, N. Lopez-Bigas, Variable interplay of UV-induced DNA damage and repair at transcription factor binding sites. Nucleic Acids Res 49, 891–901 (2021).

14. J. E. Adair, Y. Kwon, G. A. Dement, M. J. Smerdon, R. Reeves, Inhibition of nucleotide excision repair by high mobility group protein HMGA1. J Biol Chem 280, 32184–32192 (2005).

15. A. Conconi, X. Liu, L. Koriazova, E. J. Ackerman, M. J. Smerdon, Tight correlation between inhibition of DNA repair in vitro and transcription factor IIIA binding in a 5S ribosomal RNA gene. EMBO J 18, 1387–1396 (1999).

16. K. Elliott et al., Elevated pyrimidine dimer formation at distinct genomic bases underlies promoter mutation hotspots in UV-exposed cancers. PLoS Genet 14, e1007849 (2018).

17. T. Douki, J. Cadet, Individual determination of the yield of the main UV-induced dimeric pyrimidine photoproducts in DNA suggests a high mutagenicity of CC photolesions. Biochemistry 40, 2495–2501 (2001).

18. T. Douki, M. Court, S. Sauvaigo, F. Odin, J. Cadet, Formation of the main UV-induced thymine dimeric lesions within isolated and cellular DNA as measured by high performance liquid chromatography-tandem mass spectrometry. J Biol Chem 275, 11678–11685 (2000).

19. L. H. Chung, V. Murray, An extended sequence specificity for UV-induced DNA damage. J Photochem Photobiol B 178, 133–142 (2018).

20. D. S. Bryan, M. Ransom, B. Adane, K. York, J. R. Hesselberth, High resolution mapping of modified DNA nucleobases using excision repair enzymes. Genome Res 24, 1534–1542 (2014).

21. M. F. Berger et al., Compact, universal DNA microarrays to comprehensively determine transcription-factor binding site specificities. Nat Biotechnol 24, 1429–1435 (2006).

22. B. Jiang, J. S. Liu, M. L. Bulyk, Bayesian hierarchical model of protein-binding microarray k-mer data reduces noise and identifies transcription factor subclasses and preferred k-mers. Bioinformatics 29, 1390–1398 (2013).

23. C. R. Goding, H. Arnheiter, MITF-the first 25 years. Genes Dev 33, 983–1007 (2019).

24. H. M. Berman et al., The Protein Data Bank. Nucleic Acids Res 28, 235–242 (2000).

25. G. Badis et al., Diversity and complexity in DNA recognition by transcription factors. Science 324, 1720–1723 (2009).

26. C. Zhu et al., High-resolution DNA-binding specificity analysis of yeast transcription factors. Genome Res 19, 556–566 (2009).

27. L. A. Barrera et al., Survey of variation in human transcription factors reveals prevalent DNA binding changes. Science 351, 1450–1454 (2016).

28. V. Martin, J. Zhao, A. Afek, Z. Mielko, R. Gordan, QBiC-Pred: quantitative predictions of transcription factor binding changes due to sequence variants. Nucleic Acids Res 47, W127–W135 (2019).

29. M. F. Berger, M. L. Bulyk, Universal protein-binding microarrays for the comprehensive characterization of the DNA-binding specificities of transcription factors. Nat Protoc 4, 393–411 (2009).

30. F. Pedregosa et al., Scikit-learn: Machine Learning in Python. J Mach Learn Res 12, 2825–2830 (2011).

31. B. J. Frey, D. Dueck, Clustering by passing messages between data points. Science 315, 972–976 (2007).

32. K. Sugasawa, UV-DDB: a molecular machine linking DNA repair with ubiquitination. DNA Repair (Amst) 8, 969–972 (2009).

33. S. Jang et al., Damage sensor role of UV-DDB during base excision repair. Nat Struct Mol Biol 26, 695–703 (2019).

34. X. Zhang et al., Genome-wide analysis of cAMP-response element binding protein occupancy, phosphorylation, and target gene activation in human tissues. Proc Natl Acad Sci U S A 102, 4459–4464 (2005).

35. P. Mao et al., ETS transcription factors induce a unique UV damage signature that drives recurrent mutagenesis in melanoma. Nat Commun 9, 2626 (2018).

36. S. Premi et al., Genomic sites hypersensitive to ultraviolet radiation. Proc Natl Acad Sci U S A 116, 24196–24205 (2019).

37. J. I. Yeh et al., Damaged DNA induced UV-damaged DNA-binding protein (UV-DDB) dimerization and its roles in chromatinized DNA repair. Proc Natl Acad Sci U S A 109, E2737–2746 (2012).

38. Y. Takayama, D. Sahu, J. Iwahara, NMR studies of translocation of the Zif268 protein between its target DNA Sites. Biochemistry 49, 7998–8005 (2010).

39. Y. Belo et al., Unexpected implications of STAT3 acetylation revealed by genetic encoding of acetyl-lysine. Biochim Biophys Acta Gen Subj 1863, 1343–1350 (2019).

40. A. L. Stelling et al., Infrared Spectroscopic Observation of a G-C(+) Hoogsteen Base Pair in the DNA:TATA-Box Binding Protein Complex Under Solution Conditions. Angew Chem Int Ed Engl 58, 12010–12013 (2019).

41. T. Mori et al., Simultaneous establishment of monoclonal antibodies specific for either cyclobutane pyrimidine dimer or (6-4)photoproduct from the same mouse immunized with ultraviolet-irradiated DNA. Photochem Photobiol 54, 225–232 (1991).

42. J. G. Mackinnon, H. White, Some Heteroskedasticity-Consistent Covariance-Matrix Estimators with Improved Finite-Sample Properties. J Econometrics 29, 305–325 (1985).

43. J. S. Long, L. H. Ervin, Using heteroscedasticity consistent standard errors in the linear regression model. Am Stat 54, 217–224 (2000).

