## Supplementary Material for "UV irradiation remodels the specificity landscape of transcription factors"

### SUPPLEMENTARY METHODS

#### UV-Bind library designs

All DNA sequence libraries in this study were carefully designed to reduce and control the possible positions of UV pyrimidine dinucleotide photoproducts. Below are descriptions of libraries used throughout the manuscript.

*Universal UV-Bind library (Fig. 2; S2; S3; S5; S6)*

Universal libraries that contain all possible k-mer sequences are a powerful and commonly used method to determine protein binding specificities (1-3). However, traditional universal libraries contain many adjacent pyrimidine dinucleotides, which are the main "sequence substrates" for the formation of UV photoproducts (**Fig. S1A**). Thus, we designed universal UV-Bind sequences: a unique sequence library that takes advantage of the strengths of traditional universal libraries, while limiting the number of potential UV lesions in each sequence (**Fig. S1B; Table S1**). The main differences between the traditional universal library and the UV-Bind universal library are:

- The maximally compact sequences in UV-Bind are 14bp instead of 35bp in typical libraries, in order to reduce the number of pyrimidine dinucleotides in a given sequence.
- The primer used in UV-Bind is longer (35bp compared to 24bp for traditional universal libraries) and has a sequence composition that contains no pyrimidine dinucleotides to prevent dipyrimidine photoproducts from forming in the primer region.
- Additional spacer sequences between the maximally compact region, the primer and the edges. These enable a design without pyrimidine dinucleotides at the junction between the maximally compact sequence and the surrounding sequences.

This design reduces the median number of pyrimidine dinucleotides per sequence from 27 in the original universal protein binding microarray (PBM) libraries to 6. More information on the universal UV-Bind sequences can be found in **Fig. S1** which illustrates the design and **Table S1** which contains the full list of universal UV-Bind sequences.

*Increasing pyrimidine dinucleotides library (Fig. 1E; 5B-C)*

The specificity of pyrimidine dinucleotide photoproduct formation is dependent on the identify of the dinucleotide and its flanking sequence (4). We designed a library to measure how an increasing count of each pyrimidine dinucleotide would affect the binding signal of UV photoproduct antibodies (**Fig. 1**) and UV-DDB (**Fig. 5**) in multiple sequence contexts. Three sequences were designed (Sequence A: GTATGTACGCACGTGCGTACATACG, Sequence B: ACGCACATGTGTACACATGTGCGTA, Sequence C: GTACGTACGTATATATACGTACGTA), in which pyrimidine dinucleotides TT, TC, CT, and CC were inserted at positions 6, 12, and 18 from the 5' end. **Table S2** contains all the sequences in this library and their measured values.

*MITF flanking sequence library (Fig. 3A-B)*

Transcription factor MITF has been characterized to bind to the canonical E-box, CACGTG, with a preference for T and A flanks (5). We created flanking sequences of all possible variants at the (bolded) **NT**CACGTG**AN** and **NC**CACGTG**GN** positions in the sequence context of GTATGTA**NT**CACGTG**AN**TACATACG and GTATGTA**NT**CACGTG**AN**TACATACG, respectively, to examine how flanking sequence influences the effect of UV irradiation on MITF-DNA binding. Each variant was measured with a total of 20 replicates. The sequences used in the design and their median measurements (natural log transformed) can be found in **Table S4**.

*Dinucleotide variants library (Fig 3C-D; S4)*

To obtain measurements of pyrimidine dinucleotide photoproducts in variable positions relative to a binding site, libraries were designed in which a given sequence had all possible 2-bp variations across a sliding window. This design ensures that the pyrimidine dinucleotides are present at each position that is being varied. For CREB1 (**Fig. S4**) the palindromic cAMP response element site TGACGTCA was used as a binding core with a flanking context including pyrimidine dinucleotides GCATACGGC**TGACGTCA**GCCACATA and a context without any pyrimidine dinucleotides GCATACATA**TGACGTCA**TATGTATG. The flanking context without pyrimidine dinucleotides was created by modifying the 5' flank from CATACGGC to CATACATA and using the reverse complement of it on the 3' end. EGR1 used a CGCCCACGC binding core in a flanking context of ACGTATGCA**CGCCCACGC**GTATGTA and a reverse complement context of ACATAC**GCGTGGGCG**TGCATACGTA, which do not contain pyrimidine dinucleotides in the flanks. Since the EGR1 binding site could have a single nucleotide variant of CGC**A**CACGC or CGC**G**CACGC and not contain any pyrimidine dinucleotides, dinucleotide variant libraries were created for those binding cores in order to create additional non-damageable measurements for training the ordinary least squares (OLS) model. For MITF, an E-box cpre (**Fig. 3**) and an M-box core (CATGTG; **Fig. S4**) in both orientations were used in a non-damageable flanking context of GTATGTACG**CACGTG**CGTACATACG. The sequences used and their measured values can be found in **Table S4**.

*PDB variations library (Fig 5E; S2A; S5A; S7)*

Sequences from protein-DNA complexes and all of their single nucleotide variants were used in competition experiments (**Fig. 5; S7**) and the observational data on increases in UV-Bind signal after UV irradiation (**Fig. S5**). For each sequence, a subsequence of the PDB protein-DNA complex containing the binding site was adapted to the 25bp variable region of the UV-Bind library design. The following designs are adapted subsequences from the DNA sequence in protein-DNA complexes from PDB (bold: PDB sequence; regular font: added flanks):

1DH3: GCATAC**GGCTGACGTCAGCCA**CAT
5KKQ: CGCGGA**CCAGCAGGGGGCGC**ACAT
1DUX: CACATAC**ACCGGAAGTG**CACATA
2NNY: CGC**ACAGGAAGCACTTCCTGG**CAT
1PUE: CGCGC**AAAGGGGAAGTGGG**CACATA
1NKP: GCATAC**GTAGCACGTGCTAC**ACAT
3TS8: CG**CAACATGTTGGGACATGTTC**ACA
5U01: **GCGGAAATTCCCGGGAATTTCCGC**G
1HJC: CGCGC**ACTCTGTGGTTGCG**CACAT
1BG1: GCGCA**CATTTCCCGTAAATC**ATA
1TGH: GCACATACA**TATATATA**CGCACATA
1QNE: CGCGCAT**GCTATAAAAGG**CTACAT

**<u>UV-Bind experimental conditions</u>**

Binding buffers and protein concentrations are the same for each protein as described previously (6) unless noted otherwise, with MITF and UV-DDB (independent and in competition with ETS1) using the default binding buffer. For the competition assay between ETS1 and UV-DDB, 35nM of ETS1 and 250nM of UV-DDB was used. For MITF, a 175nM concentration was used in UV and non-UV conditions.

The Penta-His and Anti-GST conjugated antibodies (described in the Methods section) were used in the same conditions as the binding buffer of the protein, with a 1:20 and 1:40 ratio respectively when used. For CPD and 6-4PP measurements (**Fig. 1D-E**), antibodies were added at a 1:400 and 1:80 ratio respectively, with an addition of 0.05% Tween to the primary antibody buffer. A secondary antibody of conjugated anti-mouse 488 was added at 1.3% volume of solution.

**Data processing of k-mers for universal UV-Bind sequences**

GenePix Results (GPR) files generated by the microarray scanner for the UV-Bind experiments were processed using the normalize_array.pl script from the Seed-and-Wobble suite (2). For all datasets, unless noted otherwise, the de Bruijn file created at this step was not used. Rather, the de Bruijn sequences were extracted from the alldata.txt file and formatted so that each de Bruijn sequence was in the same position, rather than the variable positions used in the array design, to accommodate for the fixed range input required by the seed_and_wobble.pl script. The sequences were sorted by the adjusted signal, as done by default.

**Data processing of position weight matrices (PWMs)**

PWMs were generated using the seed-and-wobble.pl script from the Seed-and-Wobble suite (2) with the start and end parameters set to 1 and 14 to match the 14bp de Bruijn sequences. Because the k-mer data is at an arbitrary orientation, the orientation of the PWM models produced by seed-and-wobble.pl is also arbitrary. The lengths of the models created from seed_and_wobble.pl are a fixed length, necessitating that they are subset and aligned to compare equivalent positions. To subset the models so they represent the binding core and a flanking region of 3 bp, we took the non-UV models and calculated the core binding site region based on the top scoring position of a consensus site from literature (see below), across both orientations of the model. We then extended this range by 3 positions on either side, if possible, in order to capture the effects of flanking sequences. The core positions of the trimmed model were then compared to the UV model across all positions in both orientations. The sum of the Kullback-Leibler divergence was minimized to select the best matching position to align the models for comparison.

Binding cores used in the processing of PWMs were chosen based of literature using known consensus cores and/or structural data of TF-DNA complexes indicating positions with hydrogen bonds between the TF and DNA. For c-MYC and MITF, members of the bHLH structural family, the E-box CACGTG was used as a core and has been identified as a canonical binding site for both c-MYC (7) and MITF (5). CREB1 used the cyclic-AMP response element, TGACGTCA, for the core definition (8, 9). A binding core of CGCCCACGC was used for EGR1 based on structural data (10). For the ETS structural family members ETS1 and ELK1, a binding core of TTCC was used (11). GATA1 used a binding core of AGATAA based on structural data and the known consensus motif of WGATAR (12). STAT3 used a binding core of TCCCGGA based on structural data (13). TBP used a binding core of the TATA-box TATAAAAG (14). For P53, the binding core of a half site was used, CATG, due to the PWM motif representing a half site of the protein binding consensus (15).

**Concatenation of k-mers for UV-preferred sites**

First, 6-mers from UV-bind universal sequences were generated using the default file output from the normalize_array.pl script with the parameters “startposition” as 6 and “spotlength” as 19 for universal sequences of all 9-mers and 8-mers. An OLS with HC3 coefficients was trained on non-UV 6-mer inter-replicates using the Statsmodels python package (16). Then, 6-mers from UV conditions were compared to the OLS model. If a 6-mer was above the 99% prediction interval, it was considered a UV-preferred 6-mer (**Fig. S6B**).

The method to generate concatenated k-mers takes an initial k-mer as input and extends the sequence in one direction using the k-mer with the highest UV-preference score from the UV-preferred k-mers until the sequence can no longer be extended. Then the string is extended in the other direction until it can no longer be extended. If a k-mer is used to extend the sequence, it can no longer be used to extend the sequence. This is done for all UV-preferred k-mers using both a left, right extension and a right, left extension (**Fig. S6C)**.

This result generates a list of sequences made of concatenated k-mers of varying lengths, but the UV-bind assay only takes 25bp of variable sequence. Thus, the result of our concatenation procedure was split into two groups, one where sequences were 25bp or longer and another with sequences shorter than 25bp. For the first candidate group of sequences, those 25bp or longer, all unique 25-mers were added.

Shorter sequences were extended with a greedy method that optimizes the difference between UV and non-UV E-scores. This was done by extending sequences in an alternating left, right pattern using the k-mer with the largest difference until the sequence was 25bp. These were added to the second group of candidate sequences.

Because of the larger number of sequences and the similarity of many sequences to each other, a clustering approach was used to cluster together similar sequences and sample one sequence from each cluster to be included in the array design (**Fig. S6D**). Affinity propagation using Scikit-learn's Affinity_Propagation function was used to cluster the sequences with the random_state argument set to 0 so that the result is reproducible (17). Since many sequences were generated using a sliding window, their high similarity to each other is described using a Levenshtein distance which takes insertions and deletions into account (rather than a hamming distance for example). The distance matrix for the clustering was calculated by the minimum Levenshtein distance of both orientations: $min(lev(a,b), lev(a, reverse\_complement(b)))$.

From these clusters, the sequence with the maximum sum of UV-preference scores (**Fig. S6E**) was chosen as a representative sequence. A total of 3,200 sequences for CREB1 (2882 from the first group and 318 from the second group) and 1,884 sequences for EGR1 (1208 from the first group, 676 from the second group) were added to the library design in this manner.

**<u>Data analysis for ETS1 and UV-DDB competition</u>**

Binding measurements to the PDB library group **2NNY** (CGC**ACAGGAAGCACTTCCTGG**CAT) and **1DUX** (CACATAC**ACCGGAAGTG**CACATA) were compared due to these groups having the highest median ETS1 binding signals respectively among all groups (**Fig. S7A**). Individual measurements of sequences were binned using their distance from the diagonal with the Jenks natural breaks (18) technique for 3 bins, labeled “Weak”, “Medium” and “Strong” respectively based on the group's distance from the diagonal. For ELK1, the range of values for the binning was from the 1-99 percentile to account for outliers present in the data (see **Fig. S7B**). P-values were reported from a Jonckheere-Terpstra trend test with an alternative hypothesis for an increasing trend (19, 20) in R using the clinfun (ver. 1.1.0) package (21). Because there are more than 100 samples, the reported p-values use the built-in permutation method with the number of permutations set to $10^5$. To ensure the results can be replicated, the seed.set() method was set to 0 in R. The results for all groups can be found in **Table S5**.

### Scaling UV values in UV vs non-UV comparisons

For all CREB1, EGR1, and MITF non-k-mer data, if an analysis compared UV and non-UV conditions, the UV values were scaled to allow for a direct comparison. Using the median values for each sequence, a linear regression model using the SciPy linregress function (22) for all sequences without pyrimidine dinucleotides (UV control sequences) was trained. Using the slope and intercept from the linear regression, the values of the UV irradiated sequences were transformed so that the slope of the UV control sequences would be 1 using the formula: $(x - intercept)/slope$. The scaled values are used in figures and tables unless noted otherwise. The squared Pearson correlation coefficient and confidence interval of the scaled slope after scaling for data used in multiple figures and tables are listed below:

CREB1 data in Fig. 4, S4, and S5B; Table S4 and S5 ($R^2$ = 0.92; Slope 95% CI = 0.027)
EGR1 data in Fig. 4, S4, and S5B; Table S4 and S5 ($R^2$ = 0.91; Slope 95% CI = 0.028)
MITF data in Fig. 3, S4; Table S4 ($R^2$ = 0.89; Slope 95% CI = 0.031)

### SUPPLEMENTARY FIGURES

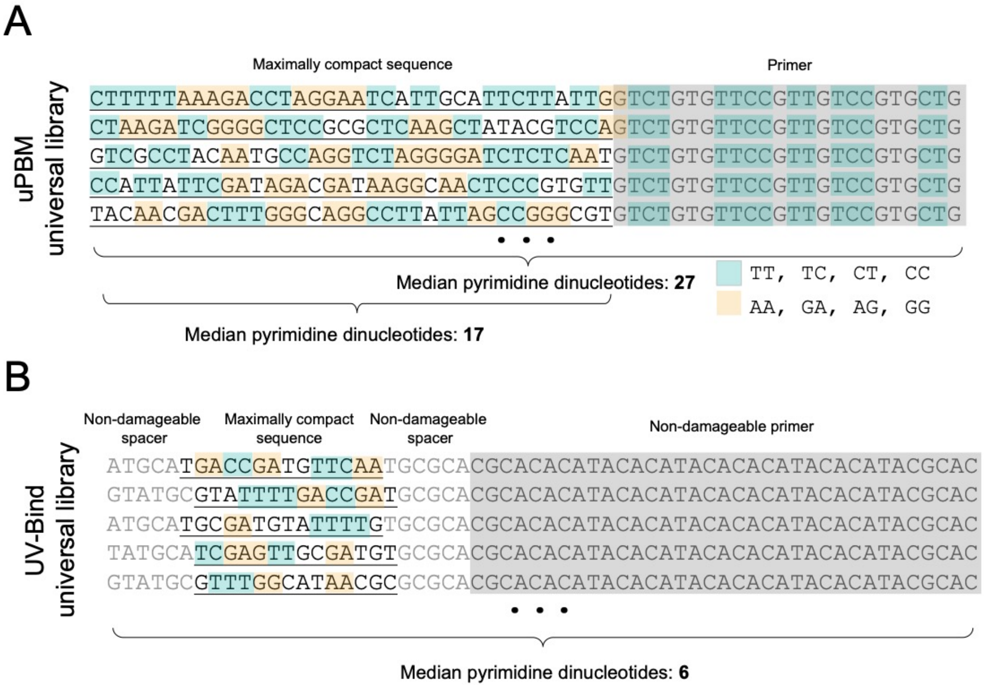

**Supplementary Figure 1:** Overview of the universal UV-Bind library design for measuring protein binding to maximally compact sequences.
**(A)** Description of the original universal protein binding microarray library. Each sequence contains part of a maximally compact sequence (or de Bruijn sequence; underlined) of 36bp and a primer (highlighted in grey) of 24bp. Positions with pyrimidine dinucleotides are highlighted in blue for the forward strand and orange for the reverse strand. Overall, there is a median value of 27 pyrimidine dinucleotides considering the whole length of the probes, and a median of 17 considering only the maximally compact sequence.
**(B)** Design overview of the maximally compact sequences in UV-Bind. The maximally compact sequence region is a 14bp subsequence flanked by non-damageable spacers (5-6bp) and followed by a non-damageable primer (35bp). It shifts 1bp in order to avoid pyrimidine dinucleotides between the sequence and the neighboring spacers. This design reduces the median number of pyrimidine dinucleotides per sequence to 6.

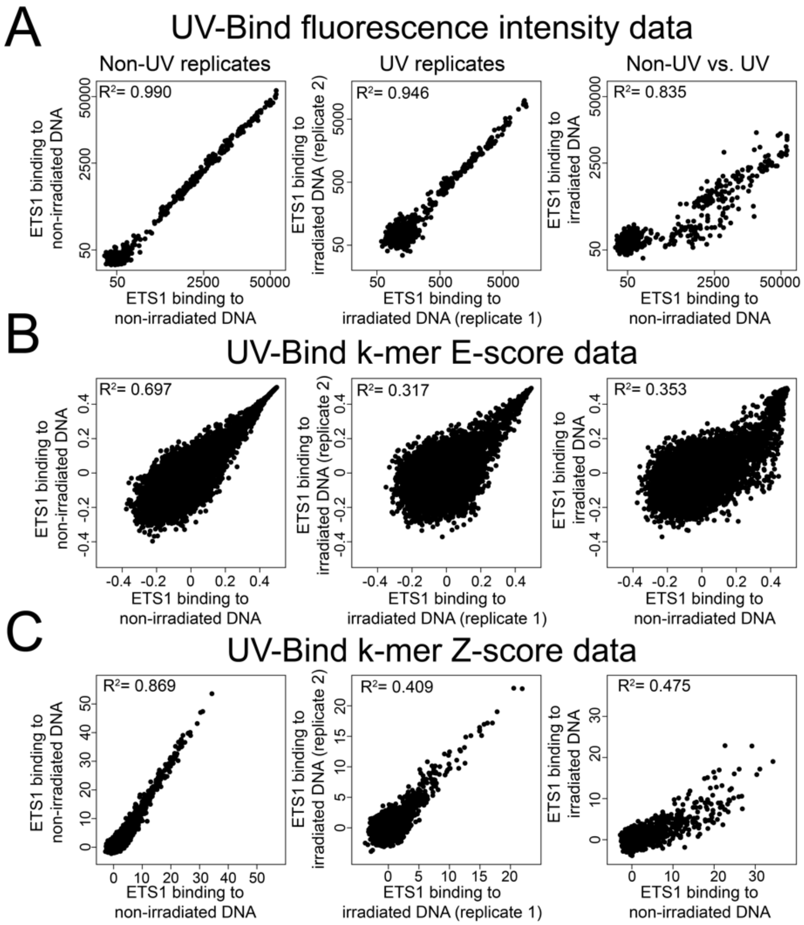

**Supplementary Figure 2:** Replicates of protein binding measurements for ETS1. Comparisons for all data are organized into non-UV replicates (left), UV replicates (middle) and a comparison among conditions (right).
**(A)** Replicates for non-universal sequences from the PDB variation library. Values represent median binding intensity.
**(B)** E-scores for replicate k-mers derived from UV-Bind measurements.
**(C)** Z-scores for replicate k-mers derived from UV-Bind measurements.

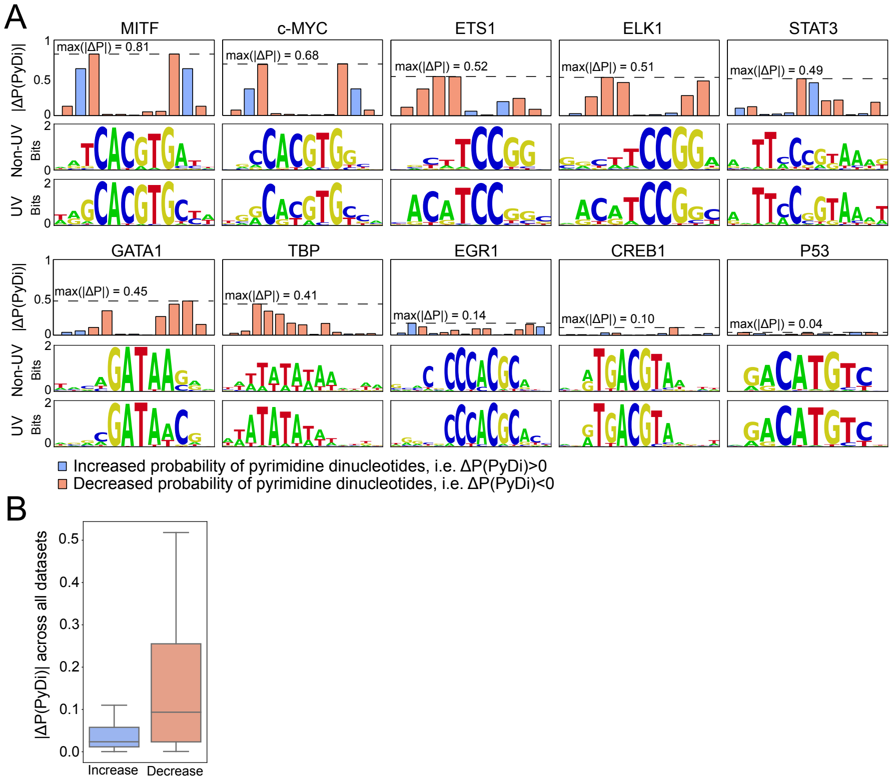

**Supplementary Figure 3:** Changes in the probabilities of pyrimidine dinucleotides due to UV irradiation, according to PWM models.
**(A)** Across 10 TFs, the absolute change in the formation any pyrimidine dinucleotide for each neighboring pair (top). A dashed line representing the largest change with the corresponding value is given. The PWM logos for the Non-UV (middle) and UV (bottom) are given, as in **Fig. 2**. The order the TFs are presented in is from the one with the largest change to the smallest.
**(B)** Comparison of the magnitude of all increases vs. all decreases. Aggregating across positions, the decreases are larger than the increases ($p = 0.0021$; Wilcoxon rank-sum).

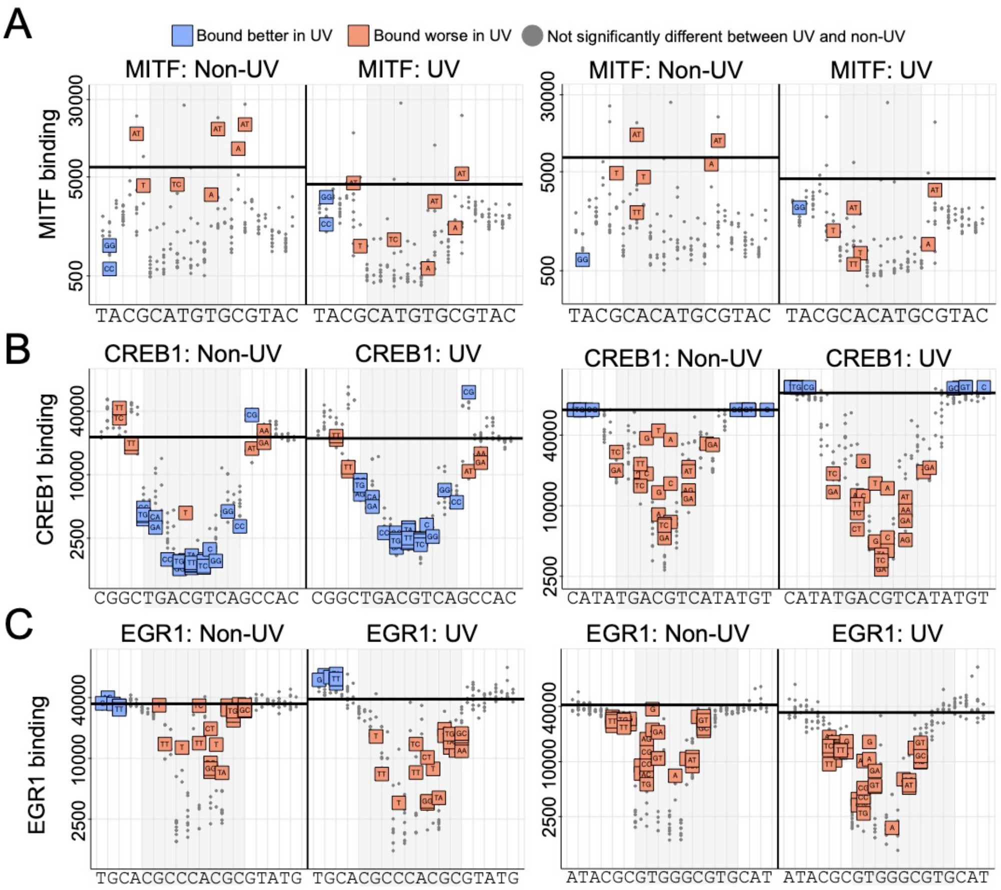

**Supplementary Figure 4:** TF binding measurements for sequence variation libraries reveal positions where UV-irradiation of DNA significantly changes TF binding. Variations with significant increases (blue) and decreases (orange) are reported as squares with the variation in the box. Non-significant changes are reported as grey dots.
(**A**) Variations for the MITF M-box CATGTG (left) and its reverse complement (right).
(**B**) Variations for the cAMP response element in a damageable (left) and non-damageable (right) flanking sequence context.
(**C**) EGR1 core sequence CGCCCACGC in non-damageable flanks (left) and its reverse complement (right).

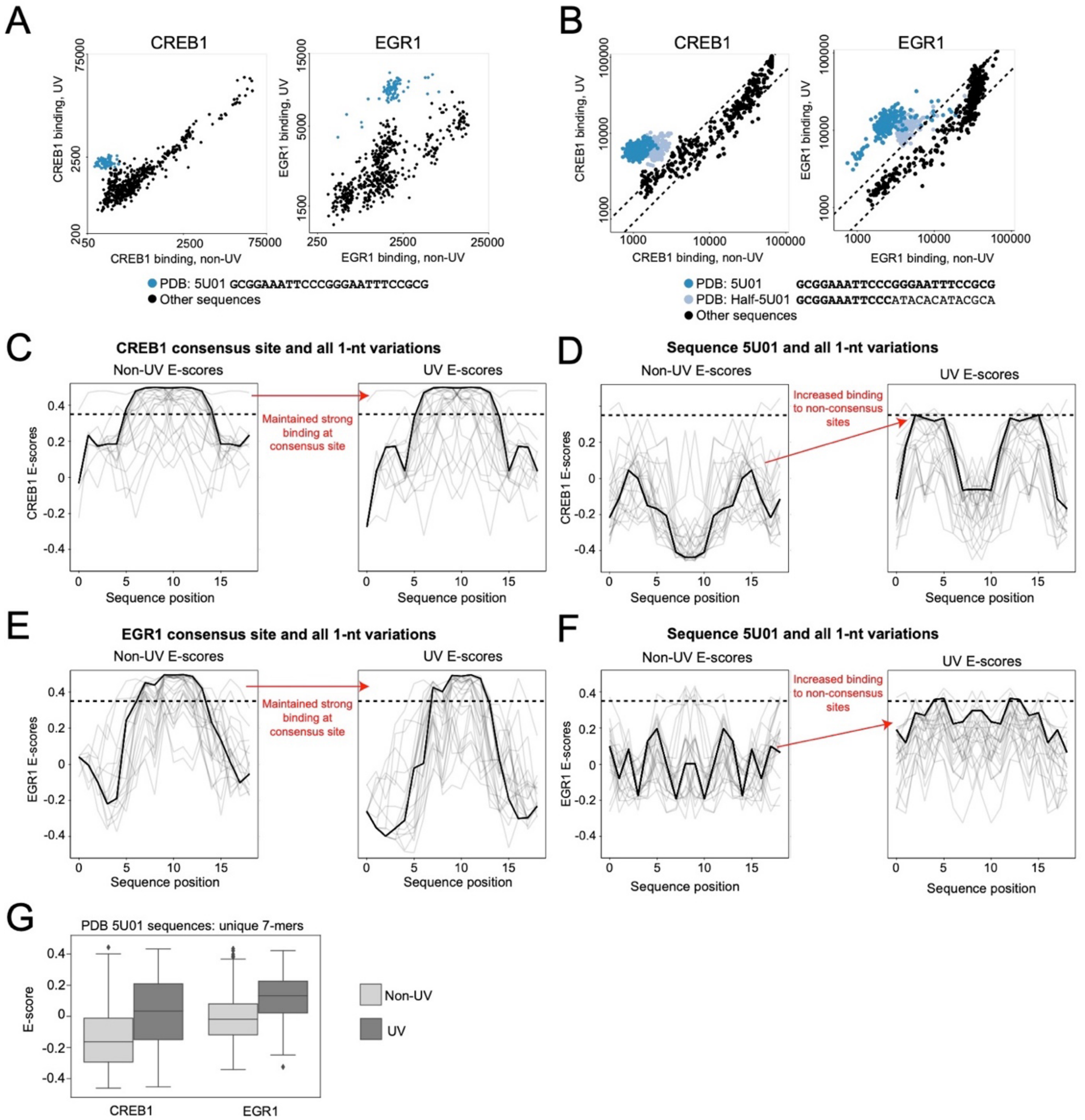

**Supplementary Figure 5:** Increase in binding of CREB1 and EGR1 to the PDB 5U01 DNA sequence and its variants after UV irradiation.

**(A)** Scatterplots of sequences in the PDB variation library for comparing non-UV and UV binding conditions for CREB1 (left) and EGR1 (right). A single nucleotide variant group based on the PDB 5U01 sequence is highlighted in blue. All the other sequences in the library are shown in black.

**(B)** Scatterplots for a follow-up experiment to (A) for a dinucleotide variant group of CREB1 (left) and EGR1 (right) sites. Dashed lines represent a 99% prediction interval from an OLS regression on probes lacking pyrimidine dinucleotides, using sequences in **Table S4**.

**(C,D)** E-score profiles for CREB1 for a library of a consensus site and 1-nt variations (C) and the 5U01 site and 1-nt variations (D). Lines are drawn with a transparency effect, so the black line in each figure represents a consensus profile among the variants while the grey lines represent variations in the profile. The dashed line represents an E-score cutoff of 0.35.

**(E, F)** Same as in (C,D) but for EGR1.

(**G**) Comparison of the unique 7-mers for the 5u01 sequence in non-UV and UV conditions.

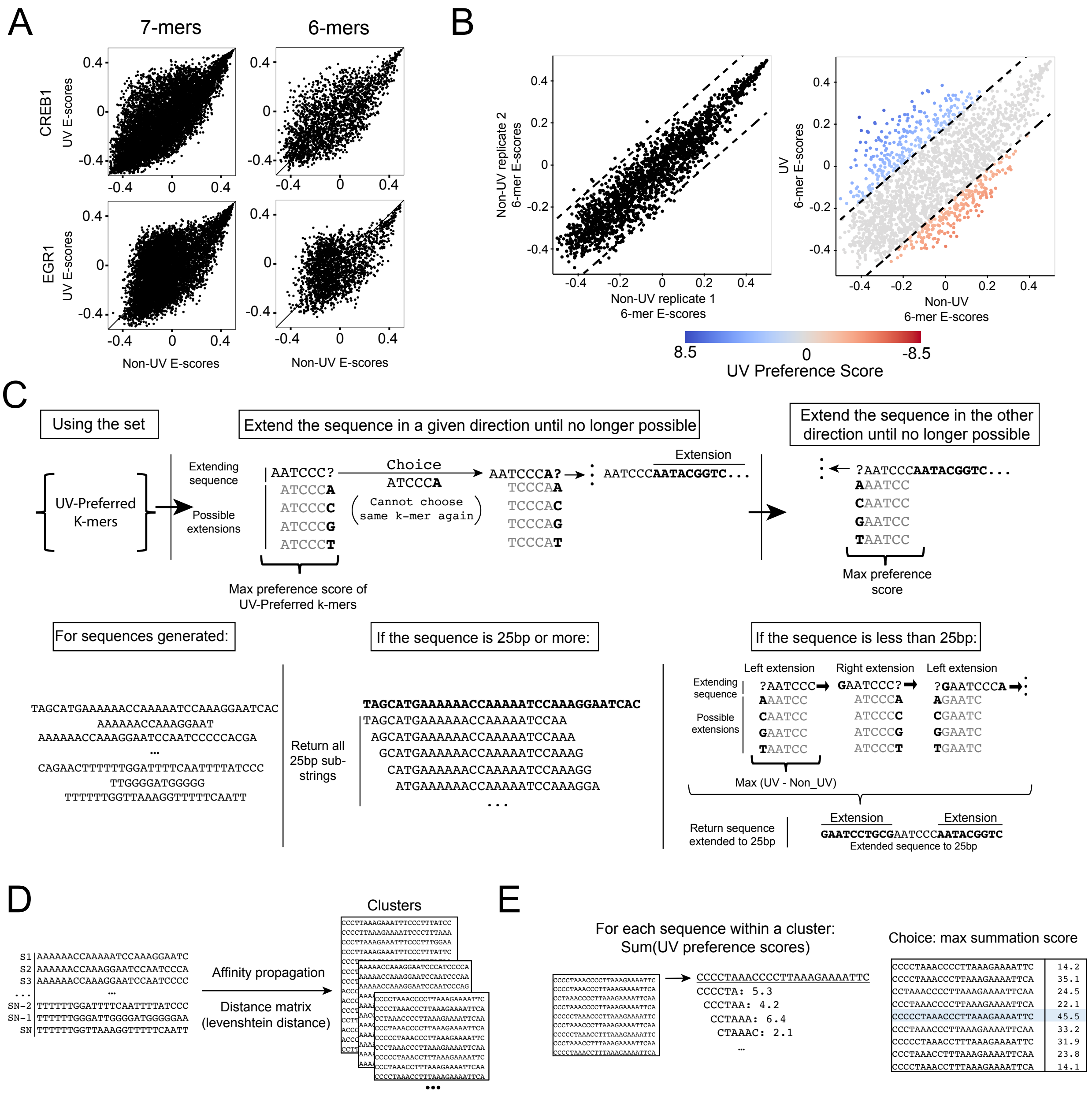

**Supplementary Figure 6:** Method for creating UV-preferred sequences by concatenating UV-preferred k-mers.
**(A)** 7-mer profiles resemble 6-mer profiles when comparing UV and non-UV conditions. Comparison of 7-mer profiles (left) to 6-mer profiles (right) for CREB1 (top) and EGR1 (bottom). For the analysis, 6-mers were used due to the availability of replicate data in non-UV and UV conditions at the time of the design.
**(B)** Classification of UV k-mers into UV-preferred (blue) and non-UV-preferred (red). An OLS model was trained on replicate 6-mers in non-UV conditions with a 99% prediction interval (left). For a comparison of non-UV vs UV conditions, k-mers above the prediction interval are considered to be UV-preferred.
**(C)** Overview of the k-mer concatenation method. K-mers were concatenated based on UV-preferred sequences and placed into 25bp contexts to fit in the sequence library.
**(D)** The concatenated k-mers were clustered using affinity propagation with the distance matrix defined by the minimum Levenshtein distance of each sequence with both orientations of the other sequence.
**(E)** The sequence with the maximum sum of UV preference scores was selected from the cluster to include in the library design.

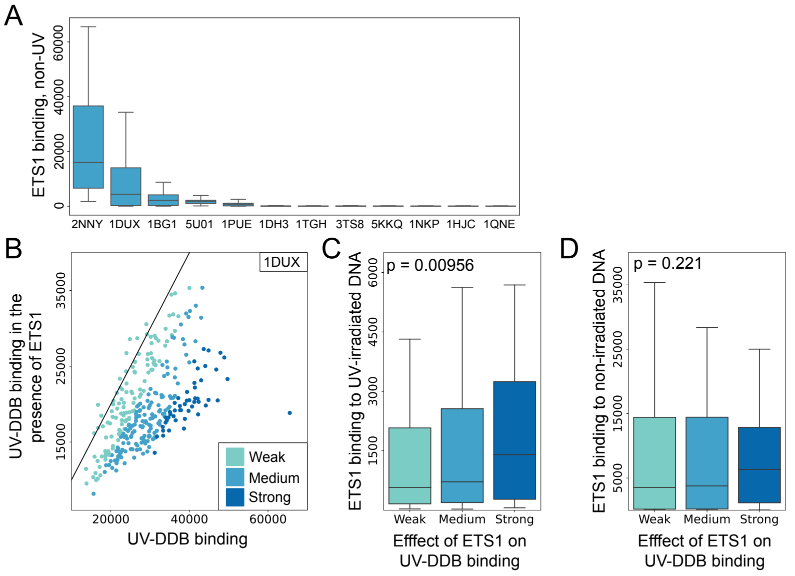

**Supplementary Figure 7:** ETS1 competes with UV-DDB for binding to UV-irradiated DNA.
**(A)** ETS1 binding to PDB sequence groups represented in our DNA libraries. Two groups, which are 1-nt variations of PDB sequences 2NNY and 1DUX, show the highest binding signals. Both sequences contain the TTCC binding core for the ETS family, which include the ETS1 TF.
**(B)** ETS1 and UV-DDB competition for 1-nt variations of a single ETS binding site, from PDB sequence 1DUX. Binning was done on the 1-99 percentile to account for outlier measurements. The scatterplot shows equidistant bins based on the 1-99 percentile with a black line representing the diagonal and points representing individual measurements of DNA sequences (the 1DUX sequence or one of its 1-nt variants).
(**C**) The ETS1 binding under UV conditions shows a consistent increasing trend as the bins move further away from the diagonal (**Table S7**).
(**D**) The ETS1 binding under non-UV conditions does not show a consistent increasing trend (**Table S7**). Trends were assessed using a Jonckheere trend test with an increasing trend as the alternative hypothesis.

### SUPPLEMENTARY TABLE CAPTIONS

Table S1. Universal UV-Bind sequence library and CPD/6-4PP data

Table S2: Measurements of CPD, 6-4PP and UV-DDB to sequences with increasing numbers of pyrimidine dinucleotides

Table S3. Universal UV-Bind data (PWMs, Z-scores, and E-scores) for the 10 TF proteins tested

Table S4: UV-Bind data for consensus binding sites with single and double nucleotide variations in the core or flanking regions

Table S5: UV-Bind data showing increased TF binding levels at non-consensus sites after UV irradiation

Table S6: UV-DDB k-mer binding data and PWM model

Table S7: Competition Data

### SUPPLEMENTARY REFERENCES

1. L. A. Barrera *et al.*, Survey of variation in human transcription factors reveals prevalent DNA binding changes. *Science* **351**, 1450-1454 (2016).
2. M. F. Berger *et al.*, Compact, universal DNA microarrays to comprehensively determine transcription-factor binding site specificities. *Nat Biotechnol* **24**, 1429-1435 (2006).
3. Z. Wang, W. He, J. Tang, F. Guo, Identification of highest-affinity binding sites of yeast transcription factor families. *Journal of Chemical Information and Modeling* **60**, 1876-1883 (2020).
4. L. H. Chung, V. Murray, An extended sequence specificity for UV-induced DNA damage. *J Photochem Photobiol B* **178**, 133-142 (2018).
5. C. R. Goding, H. Arnheiter, MITF-the first 25 years. *Genes Dev* **33**, 983-1007 (2019).
6. A. Afek *et al.*, DNA mismatches reveal conformational penalties in protein-DNA recognition. *Nature* **587**, 291-296 (2020).
7. T. K. Blackwell, L. Kretzner, E. M. Blackwood, R. N. Eisenman, H. Weintraub, Sequence-specific DNA binding by the c-Myc protein. *Science* **250**, 1149-1151 (1990).
8. M. R. Montminy, K. A. Sevarino, J. A. Wagner, G. Mandel, R. H. Goodman, Identification of a cyclic-AMP-responsive element within the rat somatostatin gene. *Proceedings of the National Academy of Sciences* **83**, 6682-6686 (1986).
9. X. Zhang *et al.*, Genome-wide analysis of cAMP-response element binding protein occupancy, phosphorylation, and target gene activation in human tissues. *Proc Natl Acad Sci U S A* **102**, 4459-4464 (2005).
10. T. B. Hamilton, F. Borel, P. J. Romaniuk, Comparison of the DNA binding characteristics of the related zinc finger proteins WT1 and EGR1. *Biochemistry* **37**, 2051-2058 (1998).
11. F. Karim *et al.*, The ETS-domain: a new DNA-binding motif that recognizes a purine-rich core DNA sequence. *Genes & development* **4**, 1451-1453 (1990).
12. J. G. Omichinski *et al.*, NMR structure of a specific DNA complex of Zn-containing DNA binding domain of GATA-1. *Science* **261**, 438-446 (1993).
13. S. Becker, B. Groner, C. W. Muller, Three-dimensional structure of the Stat3beta homodimer bound to DNA. *Nature* **394**, 145-151 (1998).
14. J. L. Kim, S. K. Burley, 1.9 A resolution refined structure of TBP recognizing the minor groove of TATAAAAG. *Nat Struct Biol* **1**, 638-653 (1994).
15. D. Menendez, A. Inga, M. A. Resnick, The expanding universe of p53 targets. *Nat Rev Cancer* **9**, 724-737 (2009).
16. S. Seabold, J. Perktold (2010) statsmodels: Econometric and statistical modeling with python. in *9th Python in Science Conference*.
17. F. Pedregosa *et al.*, Scikit-learn: Machine Learning in Python. *J Mach Learn Res* **12**, 2825-2830 (2011).
18. G. F. Jenks, The data model concept in statistical mapping. *International yearbook of cartography* **7**, 186-190 (1967).
19. A. R. Jonckheere, A distribution-free k-sample test against ordered alternatives. *Biometrika* **41**, 133-145 (1954).
20. T. J. Terpstra, The asymptotic normality and consistency of Kendall's test against trend, when ties are present in one ranking. *Indagationes Mathematicae* **14**, 327-333 (1952).
21. V. E. Seshan, M. V. E. Seshan, Package 'clinfun'. *R package clinfun* (2018).
22. P. Virtanen *et al.*, SciPy 1.0: fundamental algorithms for scientific computing in Python. *Nat Methods* **17**, 261-272 (2020).
